# Deadenylation and Decapping Factors Cooperatively Stimulate Biochemical Activities of DEAD-Box ATPase Dhh1

**DOI:** 10.64898/2026.04.28.721454

**Authors:** Gabriel A. Braun, Rakesh Kumar, Alan G. Hinnebusch, John D. Gross

## Abstract

The DEAD-box ATPase Dhh1 (DDX6 in humans) is a general activator of 5’-3’ mRNA decay that acts between the deadenylation and decapping steps of the pathway, although the exact mechanism of its action remains unclear. Dhh1 has been shown to interact with the MIF4G domain of the central scaffold protein of the Ccr4-Not deadenylase complex, Not1, as well as the decapping activator Edc3. Although structures have been published of Dhh1 in complex with Not1^MIF4G^ or an Edc3 peptide, the impact of these interactions on the catalytic cycle of Dhh1 are unknown. Here, we show Edc3 enhances ATP and RNA binding by Dhh1, whereas Not1^MIF4G^ promotes the catalytic step of ATP hydrolysis. Additionally, the modulation of Dhh1 activity by Edc3 requires a more extensive set of interaction motifs and interfaces than was previously recognized. While the effect of either Not1^MIF4G^ or Edc3 on the ATPase activity of Dhh1 is modest, together both proteins increase Dhh1 activity over 200-fold. The fact that Dhh1 biochemical activity is cooperatively tuned by deadenylation and decapping factors suggests that Dhh1 may coordinate deadenylation and decapping in the 5’-3’ mRNA decay pathway through changes in its ATP-coupled RNA binding affinity during its catalytic cycle.

**Graphical Abstract:** 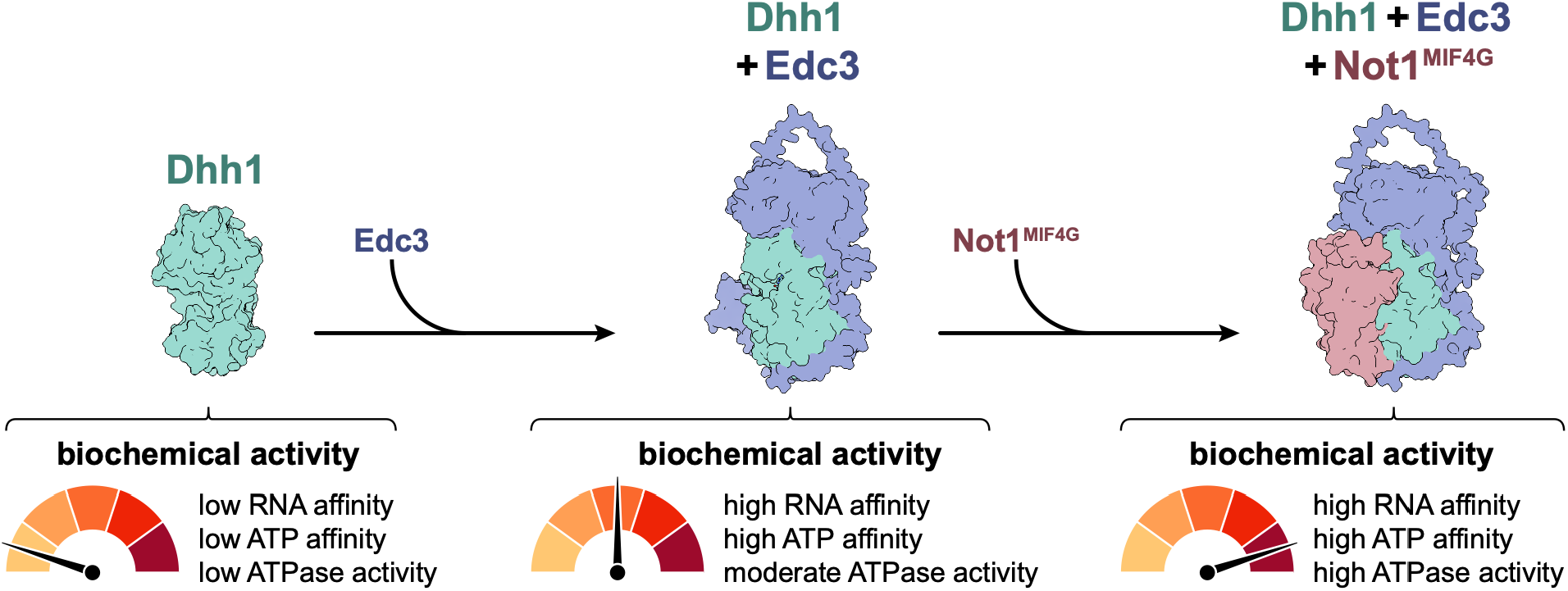

## INTRODUCTION

RNA degradation plays a critical role in post-transcriptional gene regulation, allowing for control over what transcripts are present and available for translation. The steady-state half-lives of different transcripts can vary by several orders of magnitude, and the stability of a given transcript can shift rapidly, particularly during cell-fate decisions and in response to environmental cues (1–3). The 5’-3’ mRNA decay pathway is one of the two primary means of generalized cytosolic RNA degradation and plays a key role in cell proliferation, immune response, and early organismal development, among other processes (3). This pathway begins with deadenylation, in which the mRNA 3’ poly(A) tail is shortened by two conserved deadenylases, one of which is the megadalton Ccr4-Not complex (4, 5). Following deadenylation, the transcript undergoes decapping, in which the 5’ 7-methylguanosine cap is removed by the conserved Dcp1/Dcp2 decapping complex, aided by a number of specific decapping activators (6). The exposed 5’ end of the mRNA is then vulnerable to processive 5’-3’ exonucleolysis. In addition to the nucleases and hydrolases that carry out these successive steps, many additional enzymes and cofactors play a role in 5’-3’ decay, contributing both redundancy and specificity to the pathway.

Yeast Dhh1 (DDX6/Rck/p54 in humans, Me31B in *Drosophila,* also known as Ste13 *in S. pombe*) is a conserved general activator of 5’-3’ mRNA decay and is responsible for the decay of over 1000 transcripts (7–10). Dhh1 acts between the deadenylation and decapping steps of the pathway, with its deletion resulting in accumulation of deadenylated, capped RNA (7). In addition to its role in 5’-3’ decay, Dhh1 is also involved both in codon optimality-linked decay and microRNA-mediated decay (11, 12). Beyond RNA decay, Dhh1 also contributes to both microRNA-mediated and general translational repression (11, 13–17). Dhh1 is also critical for P-body formation and maintenance, while also playing a role in stress granule formation and modulating interactions between P-bodies and stress granules (18–20). At a cellular and organismal level, Dhh1 and its orthologs are involved in a wide range of processes, including stress response, adipogenesis, progenitor cell function, cell cycle regulation, maternal mRNA storage, and oocyte development; additionally, they have been implicated in both viral infection and oncogenesis (21–24).

Dhh1 is part of the DEAD-box protein (DBP) family, which plays a key role in virtually every step of the RNA lifecycle (25, 26). DBPs are characterized by a set of nine conserved sequence motifs and share core biochemical functions, including RNA binding, ATP binding, and ATP hydrolysis (27). These canonical functions have been shown to be important for Dhh1 function *in vivo*, with mutations that perturb any of these core activities leading to deficiencies in Dhh1-dependent RNA decay, translational repression, and P-body regulation, as well as defects in cell growth and organismal development (19, 28–30). Additionally, Dhh1 has been shown to interact with over a dozen other proteins involved in 5’-3’ mRNA decay, with crystal structures solved for the binary interactions between Dhh1 and Not1 (the core scaffolding protein of the Ccr4-Not complex), and peptides from the decapping activators Edc3, Pat1, and LSM14A (Trailer Hitch/Tral in *Drosophila*, Scd6 in yeast) (31–34). In budding yeast, Edc3 and Scd6 have been shown to act redundantly to enhance the decay of a substantial fraction of total mRNAs by enhancing the association of Dhh1 with the decapping enzyme Dcp2 (14, 35).

Despite the genetic evidence that RNA binding and ATP hydrolysis are critical to the many *in vivo* functions of Dhh1, previous work has shown that the protein displays atypical *in vitro* activity for a DBP. Whereas RNA binding for most DBPs is highly coupled to nucleotide binding, Dhh1 binds RNA with equal affinity in the presence and absence of nucleotide (29). Moreover, Dhh1 has been reported to have very low intrinsic ATPase activity, likely due to autoinhibitory interdomain interactions (28, 29, 32, 33). Thus, while the *in vivo* functional importance of Dhh1 has been well established, the discrepancies between its *in vivo* and *in vitro* behavior have remained unresolved and have been a barrier to understanding the role of Dhh1 in its various biological functions.

DBP activity is often tuned by interactions with regulatory protein cofactors, which can variously cause structural rearrangement, enzymatic inhibition or stimulation, and changes to RNA affinity (27). For example, HEAT repeat-containing proteins can facilitate DBP ATPase activity through allosteric stabilization of the tandem RecA domains that form a composite active site for ATP (36–38). Consistent with this, the Not1 MIF4G domain (named for its homology to the middle domain of eIF4G) has previously been reported to stimulate the ATPase activity of Dhh1, although this stimulatory effect is relatively weak (19, 32). This suggests that previously annotated but functionally uncharacterized interactions between Dhh1 and other proteins may be key to resolving the discrepancy between the *in vivo* and *in vitro* behavior of Dhh1.

Here, we investigate using *S. pombe* proteins how both the decapping activator Edc3 and the core deadenylase subunit Not1 modulate Dhh1 activity and perhaps contribute to 5’-3’ mRNA decay. We find that Edc3 both increases the ATPase activity of Dhh1 and enhances the affinity of Dhh1 for RNA in a nucleotide-dependent manner, likely by remodeling Dhh1 to prime it for binding both ATP and RNA. Furthermore, we show that this functional modulation requires extensive multi-domain interactions between Dhh1 and Edc3, which were previously thought to interact primarily via a minimal, localized set of short motifs. We also find that, while the MIF4G domain of Not1 stimulates Dhh1 activity, it does so by a different mechanism than does Edc3. Further, we show that Edc3 and Not1^MIF4G^ not only independently increase Dhh1 activity but also combine to synergistically stimulate ATP hydrolysis. We propose a model in which Not1 and Edc3 tune the RNA binding and ATPase activity of Dhh1, thus linking the 3’ deadenylase machinery with the 5’ decapping complex to coordinate 5’-3’ mRNA decay.

## MATERIALS AND METHODS

### Protein Expression & Purification

Details on protein constructs, including amino acid boundaries, mutations, purification tags, and expression vectors are in **Table S1**.

*S. pombe* Dhh1 was expressed in *E. coli* BL21(DE3) (New England Biolabs) cultured in LB medium. Cultures were grown at 37 °C until OD_600_ = 0.5-0.8, then transferred to 4 °C for 20 minutes before being induced with 1 mM IPTG and grown for 16 hours at 18 °C. Cells were harvested by centrifugation (4,000 *g* at 4 °C for 20 minutes), resuspended in lysis buffer (20 mM HEPES, pH 7.5, 500 mM NaCl, 20 mM imidazole, 5 mM β-mercaptoethanol (BME)) supplemented with lysozyme and protease inhibitor cocktail (Roche), lysed by sonication, and centrifuged (14,500 *g* at 4 °C for 45 minutes) to pellet cell debris. The clarified lysate was loaded onto a HisTrap column (Cytiva), washed with 15 column volumes (CVs) of lysis buffer, and eluted in 5-10 CVs elution buffer (20 mM HEPES, pH 7.5, 250 mM NaCl, 250 mM imidazole, 5 mM BME). TEV protease was added to the eluate and incubated overnight at 4 °C to remove the His_6_ tag. The TEV digestion mixture was loaded onto a HiTrap heparin column (Cytiva), washed with 10 CV low salt buffer (25 mM HEPES, pH 7.5, 100 mM NaCl, 2 mM DTT), and eluted over a 10 CV gradient of low salt buffer to high salt buffer (25 mM HEPES, pH 7.5, 1 M NaCl, 2 mM DTT). Proteins were further purified by size-exclusion chromatography (SEC) using a Superdex 200 16/600 column (GE Healthcare), eluted via isocratic flow in buffer containing 25 mM HEPES, pH 7.5, 150 mM NaCl, and 1 mM DTT. Purity was assessed by SDS-PAGE and pure fractions were combined, concentrated, flash frozen in liquid nitrogen, and stored at -80 °C. Dhh1^DQAD^ was purified identically to wild-type.

*S. pombe* Edc3 was expressed as described for Dhh1. His_6_ tag purification was performed as for Dhh1, with the following changes: Lysis buffer was 50 mM HEPES, pH 7.5, 400 mM NaCl, 10 mM imidazole, 0.1% TritonX-100, and 5 mM BME, supplemented with lysozyme and protease inhibitor cocktail (Roche). Once loaded on the HisTrap column, the protein was washed with 10 CV each of high salt buffer (25 mM HEPES, pH 7.5, 400 mM NaCl, 10 mM imidazole, 5 mM BME) and low salt buffer (25 mM HEPES, pH 7.5, 100 mM NaCl, 10 mM imidazole, 2 mM BME). HisTrap Elution buffer was 25 mM HEPES, pH 7.5, 100 mM NaCl, 250 mM imidazole, and 2mM BME. TEV cleavage, heparin purification and SEC were performed as described above. Edc3^ΔLsm^, Edc3^ΔYjeF-N^, Edc3^ADA^, Edc3^HN-mut^, Edc3^YjeF-mut^, and Edc3^HN/YjeF-mut^ were all purified identically to wild-type. The Edc3^FDF^ peptide was purchased from Peptide 2.0.

*S. pombe* Not1^MIF4G^ was expressed as described for Dhh1 and induced with 0.5 mM IPTG. Cells were resuspended in lysis buffer (50 mM HEPES, pH 7.5, 300 mM NaCl, 2 mM EDTA, 1 mM DTT) supplemented with lysozyme and protease inhibitor cocktail (Roche) and lysed as described for Dhh1. The clarified lysate was loaded onto an MBPTrap column (Cytiva), washed with 15 CV wash buffer (20 mM HEPES, pH 7.5, 200 mM NaCl, 1 mM EDTA, 1 mM DTT), and eluted in 5-10 CV wash buffer supplemented with 10 mM maltose. PreScission protease was added to the eluate and incubated overnight at 4 °C to remove the MBP tag. The cleavage reaction was diluted with 20 mM HEPES, pH 7.5 to reach a final NaCl concentration of 100 mM and loaded on a HiTrap Q column (Cytiva), washed with 10 CV low salt buffer, and eluted over a 10 CV gradient of low salt buffer to high salt buffer (both buffers identical to those used for heparin purification). SEC was performed as described for Dhh1.

### ATPase Assays

Dhh1 ATPase activity was assessed by γ-^32^P ATPase assays. Reactions were carried out in 25 mM HEPES, pH 7.5, 150 mM NaCl, 5 mM MgCl_2_, and 1 mM DTT. Single-turnover reactions (with enzyme in excess of substrate) contained 1 nM γ-^32^P ATP; multiple turnover reactions (with substrate in excess of enzyme) contained 1 mM total ATP, doped with trace amounts of γ-^32^P ATP. Experiments testing a single enzyme concentration contained 5 µM Dhh1 and (as relevant) 15 µM Edc3, 15 µM Not1^MIF4G^, and 100 µM U_10_ RNA. Concentration series contained 200 µM U_10_ RNA and (as relevant) 100 µM Edc3 and 100 µM Not1^MIF4G^. All reactions contained RNA, unless specifically noted. All ATPase assays were performed in triplicate.

Reaction mixtures were preincubated for 10 minutes before being initiated by addition of ATP and MgCl_2_. Aliquots were taken at appropriate and quenched in 2x stop solution (50 mM HEPES, pH 7.5, 100 mM EDTA, 3% SDS). Inorganic phosphate was separated from ATP by thin-layer chromatography using PEI-cellulose plates run in 0.5 M LiCl, 1 M formic acid. Plates were exposed to a phosphor screen and imaged on a Typhoon Imager (GE Life Sciences). The fraction hydrolyzed ATP was quantified using ImageJ.

Observed rates (*k*_obs_) for single-turnover kinetics were determined in GraphPad Prism (v.11) by fitting data to the following equation:

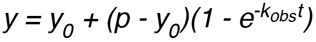

where *y_0_* is the initial fraction free P_i_, and *p* is the fraction ATP hydrolyzed at the plateau; *y*_0_ was typically set to 0 and data were fit constrained to a common *p*. Initial velocity (v0) for multiple-turnover kinetics was determined by linear regression of data points for which f_Pi_ < 0.2. Concentration series were fit to the following equation:

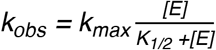

where *[E]* is the concentration of Dhh1, *k_max_* is the saturating rate constant, and *K_1/2_* is the enzyme concentration of half-maximal enzymatic activity.

### Mant-ATP Binding

Nucleotide binding assays were carried out in 25 mM HEPES, pH 7.5, 150 mM NaCl, 5 mM MgCl_2_, and 1 mM DTT. Dhh1 and, when applicable, a two-fold molar excess of Edc3 were serially diluted into buffer containing 0.5 µM methylanthraniloyl-labelled ATP (Jena Biosciences) and incubated 20 minutes. All mant-ATP binding experiments were performed in triplicate. Fluorescence was measured using a SpectraMax iD5 plate reader (Molecular Devices), with excitation at 280 nm and emission measured at 450 nm. Fluorescence data were fit using GraphPad Prism to the following equation:

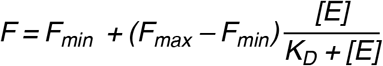

where *F_min_* and *F_max_* are the minimum and maximum fluorescence plateau values, respectively, *[E]* is the concentration of Dhh1, and *K_D_* is the dissociation constant. Because Dhh1, in the absence of Edc3, binds mant-ATP sufficiently weakly that *F_max_* could not be reliably determined, the data sets with and without Edc3 were fit together to a shared *F_max_*.

### Florescence Polarization

Fluorescence polarization (FP) assays were carried out in 25 mM HEPES, pH 7.5, 150 mM NaCl, 5 mM MgCl_2_ and 1 mM DTT. When applicable, 1 mM nucleotides (ATP or ADP) or nucleotide analogues (ADP-BeF_x_ or ADP-AlF_4_) were added to the buffer. ADP-BeF_x_ and ADP-AlF_4_ were made as previously described (39), by combining ADP with a five-fold molar excess of the metal fluoride and a 25-fold molar excess of NaF. When applicable, Edc3 and Not1^MIF4G^ were added in at least two-fold molar excess of the highest Dhh1 concentration used in a given experiment. All FP experiments were performed in triplicate.

Dhh1 was serially diluted into buffer containing 10 nM 5’ fluorescein-labeled U_20_ RNA (Integrated DNA Technologies) and incubated 20 minutes. Fluorescence polarization was measured using a SpectraMax iD5 plate reader (Molecular Devices), with excitation at 485 nm and emission measured at 535 nm. Background signal was subtracted from polarization data, which was then normalized and fit using GraphPad Prism to the following equation:

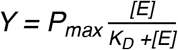

where *P_max_* is the maximal polarization, *[E]* is the concentration of Dhh1, and *K_D_* is the dissociation constant.

### Mass Photometry

Mass photometry (MP) experiments were performed with a Refeyn OneMP (Refeyn Ltd.), calibrated with BSA, apoferritin, and ADH. Samples were prepared in 25 mM HEPES, pH 7.5, 150 mM NaCl, 5 mM MgCl_2_, and 1 mM DTT. Samples were prepared in equimolar 1 µM mixtures. Ternary Dhh1/Edc3/Not1^MIF4G^ samples additionally included 1 mM ADP-BeF_x_ and 5 µM U_10_ RNA. For crosslinked samples, 0.01% glutaraldehyde was added to the prepared sample, incubated for 30 seconds, then quenched with 100 mM Tris. For each sample, 15 µL of buffer was added to the well; 1 µL of sample was diluted into the buffer; the protein concentration was adjusted by adding more sample or buffer to obtain sufficient detected events without overcrowding. Analysis was performed in DiscoverMP (Refeyn Ltd.) with the default settings.

### Protein Structure Predictions

Structural models were predicted using AlphaFold 3 (40). Models with the highest confidence were used for further analysis and visualization in UCSF ChimeraX (41).

### Cell Spotting Growth Assays

Details on plasmids, including mutations, epitope tags, and vectors are in **Table S2**.

Empty vector or Myc-tagged *EDC3* plasmids (either wild-type or mutant) were transformed into a the *scd6Δedc3Δ* double deletion *S. cerevisiae* strain FZY858 (*MATa ade2-1 ura3-1 his3-11,15 trp1-1 leu2-3,112 can1-100 scd6Δ::hphMX4 edc3Δ::kanMX4*) (14). Cells were grown to mid-logarithmic phase at 30 °C in synthetic complete medium lacking leucine (SC–Leucine). Cultures were diluted to OD_600_ = 1.0 and 10-fold serial dilutions were spotted on either SC–Leucine medium or the same medium containing 3% glycerol and 2% ethanol in place of glucose and incubated at 30 °C or 37 °C for two days before imaging.

### Statistical Analyses

All statistical analysis was performed in GraphPad Prism (v.11). Specifics of tests used are provided in figure legends. *P*-values were denoted as follows: *p* > 0.05 considered not significant (ns), \**p* ≤ 0.05, \*\**p* ≤ 0.01, \*\*\**p* ≤ 0.001, and \*\*\*\**p* ≤ 0.0001.

## RESULTS

### Edc3 is an activator of Dhh1

While Edc3 is known to interact with Dhh1 both *in vivo* and *in vitro* (33, 42, 43), the effects of this interaction on Dhh1 enzymatic activity have remained unknown, with previous studies noting only that Dhh1 ATPase activity is virtually unmeasurable (33, 34). Since these studies used Dhh1 constructs lacking its N- and C-terminal intrinsically disordered regions and various short (30- to 80-mer) Edc3 constructs, there are potentially functionally important interactions present in the full-length proteins that would not have been observed. Accordingly, we performed single and multiple turnover ATPase assays with full-length *S. pombe* Edc3 and Dhh1 proteins in the presence of excess U_10_ RNA, which typically strongly stimulates the enzymatic activity of DBPs. (Like most DBPs, Dhh1 shows minimal RNA sequence specificity; as such, polyuridylic acid has been adopted as a standard *in vitro* substrate (19, 25, 27, 29, 32, 44)).

In single-turnover conditions, we find that Edc3 stimulates the ATPase activity of Dhh1, with a five-fold increase in rate, indicating that Edc3 directly enhances either the binding or the hydrolysis of ATP by Dhh1 (**Fig. S1a-b**, **Table S3**). Under multiple-turnover conditions, Edc3 stimulates Dhh1 activity 25-fold (**Fig. S1c-d**, **Table S4**). The fact that Edc3 stimulates Dhh1 to a greater extent under multiple-turnover conditions suggests that Edc3 also accelerates a post-hydrolysis step of the Dhh1 catalytic cycle, possibly by alleviating ADP product inhibition. Consistent with previous reports, we find that Dhh1 alone exhibits very low enzymatic activity, with an observed rate under single-turnover conditions of under 0.05 h^-1^ and barely observable activity under multiple turnover conditions, even on hours-long timescales (**Fig. S1**, **Tables S3-4**).

To define the mechanism by which Edc3 stimulates Dhh1 activity, single-turnover kinetic experiments were carried out at a range of enzyme concentrations for Dhh1 both alone and in the presence of saturating amounts of Edc3 (**Fig. 1a-b**). Fitting the observed rate, *k*_obs_, as a function of enzyme concentration allows for determination of the maximal rate, *k*_max_, and the concentration at which half-maximal activation is reached, K_1/2_ (these values are analogous to the Michaelis-Menten parameters *v*_max_ and K_M_, respectively). Dhh1 alone has a maximal rate of under one per hour, with a high double-digit micromolar K_1/2_ (**Table S5**). This K_1/2_ value likely reflects a low affinity for substrate (ATP), consistent with moderate-to-low affinities for ATP among DBPs generally (25, 45, 46). In the presence of Edc3, the maximal rate of Dhh1 is not significantly increased but the K_1/2_ is reduced almost 50-fold (**Fig. 1b**, **Table S5**). We conclude that Edc3 stimulates Dhh1 by making the enzyme more active at low concentrations, likely by enhancing its affinity for ATP. To examine the latter possibility, we tested the affinity of Dhh1 for ATP labeled with a fluorescent methylanthraniloyl (mant) moiety in the absence and presence of Edc3 (**Fig. 1c-d**). Indeed, we find that the affinity of Dhh1 for mant-ATP is significantly increased in the presence of Edc3, consistent with Edc3 directly increasing the affinity of Dhh1 for ATP (**Table S6**). (The fact that the K_1/2_ for ATPase activity is slightly lower than the K_D_ for mant-ATP binding is likely caused by the mant moiety, which has previously been reported to modestly decrease affinity for nucleotide in other systems (47, 48).)

**Figure 1.**
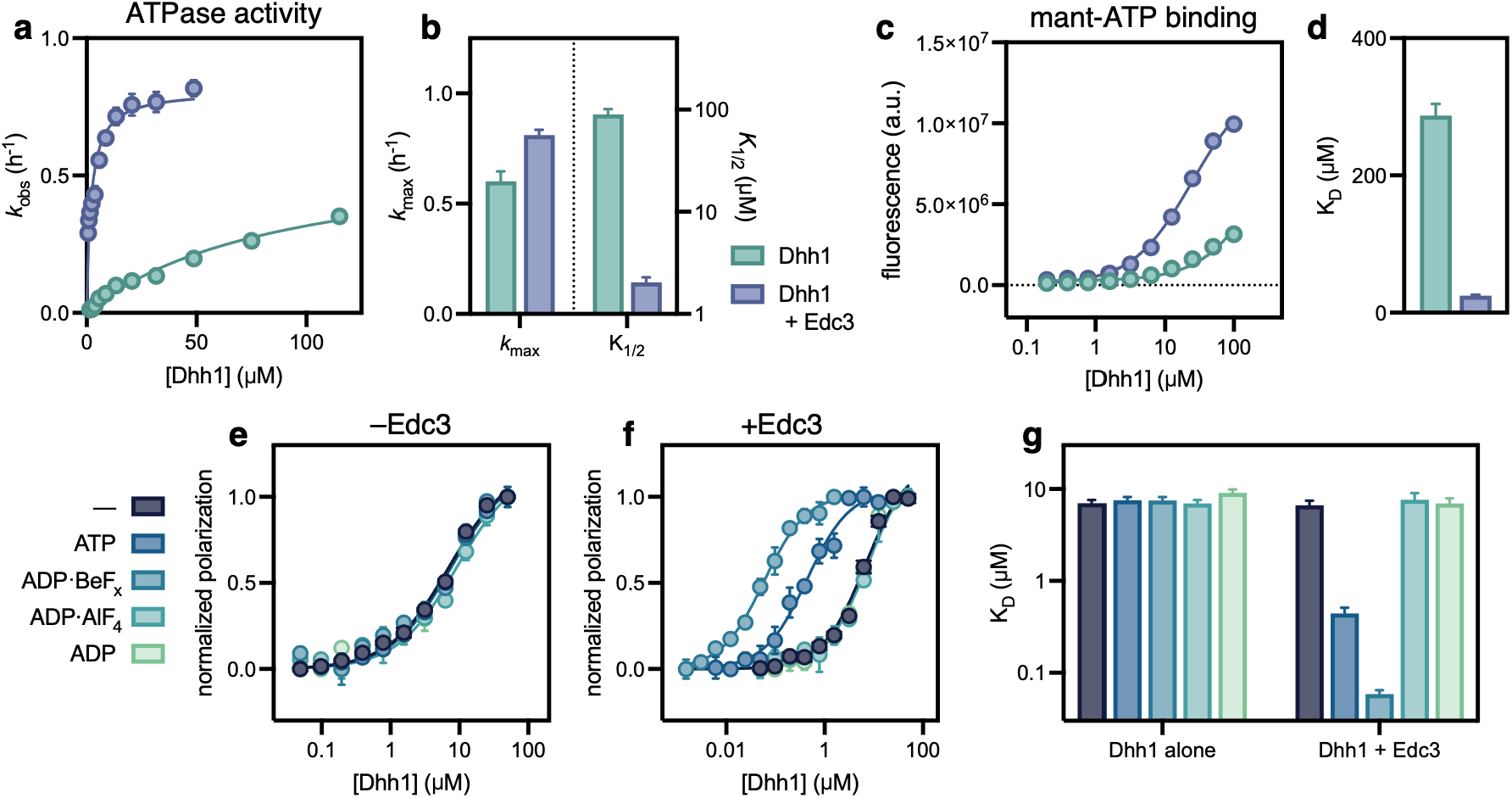
Edc3 stimulates Dhh1 ATPase activity and RNA binding. **a.** Rate of single-turnover Dhh1 ATPase activity as a function of Dhh1 concentration in the absence (green) and presence (blue) of saturating amounts of Edc3. All reactions contain saturating amounts of U_10_ RNA. Each point represents fitted rate and standard deviation from n = 3 technical replicates. **b.** Values for *k*_max_ and K_1/2_ of Dhh1 activity in the absence and presence of Edc3, derived from single-turnover kinetic data shown in panel (a), with error bars showing standard deviation. **c.** Fluorescence binding assay showing binding of Dhh1 to mant-ATP in the absence (green) and presence (blue) of saturating amounts of Edc3. Data points show mean and standard deviation of n = 3 technical replicates. **d.** Fitted K_D_ values for data shown in panel (c), with error bars showing standard deviation. **e-g.** Fluorescence polarization showing Dhh1 binding to a fluorescein-labeled U_20_ RNA in the **(e)** absence and **(f)** presence of saturating amounts of Edc3 in the absence of nucleotide or with various nucleotides and nucleotide analogs. Data points show mean and standard deviation of n = 3 technical replicates. **g.** Fitted K_D_ values for data shown in panels (e) and (f), with error bars showing standard deviation. See **Table S5** for *k*_max_ and K_1/2_ values shown in panel (b) and **Tables S6-7** for K_D_ values shown in panels (d) and (g), respectively.

Given that Edc3 enhances the affinity of Dhh1 for ATP, we next asked whether Edc3 likewise enhances the binding of RNA by Dhh1. Most DBPs display strong coupling between ATP and RNA binding, with the affinity for RNA changing by as much as several orders of magnitude depending on the nucleotide state (27, 46, 49). We used fluorescence polarization (FP) to follow binding of Dhh1 to a fluorescein-labeled U_20_ RNA in the absence of nucleotide or in the presence of ATP, ADP, ADP-beryllium fluoride (ADP-BeF_x_), or ADP-aluminum fluoride (ADP-AlF_4_) (**Fig. 1e-g**). (The nucleotide analogs ADP-BeF_x_ and ADP-AlF_4_ have been used previously to study the ATP usage of DBPs (39, 50).)

In the presence of either ATP or ADP-BeF_x_ (an ATP pre-hydrolysis structural mimic), Edc3 enhances the affinity of Dhh1 for RNA by 10- and 100-fold, respectively (**Fig. 1e-g**, **Table S7**). In contrast, in the absence of nucleotide or in the presence of either ADP or ADP-AlF_4_ (an ATP transition state mimic), Edc3 has no effect on the affinity of Dhh1 for RNA. The fact that we observe lower affinity for RNA in the presence of ATP than with ADP-BeF_x_ is likely due to the fact that, with ATP, a population of Dhh1 has hydrolyzed ATP and is in the low RNA-affinity, ADP-bound state (39). Taken together, these data suggest that the enhancement of Dhh1 affinity for RNA by Edc3 requires the ATP bound, pre-hydrolysis state of Dhh1 and, moreover, that ATP hydrolysis by Dhh1 would result in subsequent loss of RNA affinity. Consistent with previous reports, we find that in the absence of Edc3, Dhh1 displays no nucleotide dependence for RNA binding, contrary to most DBPs (**Fig. 1e**,**g**, **Table S7**) (29, 33, 51). These results show that, beyond simply enhancing the affinity of Dhh1 for RNA, Edc3 couples ATP and RNA binding, sensitizing Dhh1 affinity for RNA to nucleotide state as is typical for DBPs.

### Multiple domains of Edc3 are necessary for activation of Dhh1

Edc3 contains two structured domains connected by a long intrinsically disordered region (IDR) (**Fig. 2a**). The N-terminal Lsm domain interacts with the Dcp1/Dcp2 decapping complex (52, 53); the C-terminal YjeF-N domain mediates homodimerization of Edc3 (54); and the interdomain IDR contains Phe-Asp-Phe (FDF), Phe-Asp-Lys (FDK), and tryptophan (Trp) motifs that have previously been shown to be involved in the interaction with Dhh1 (33, 34).

**Figure 2.**
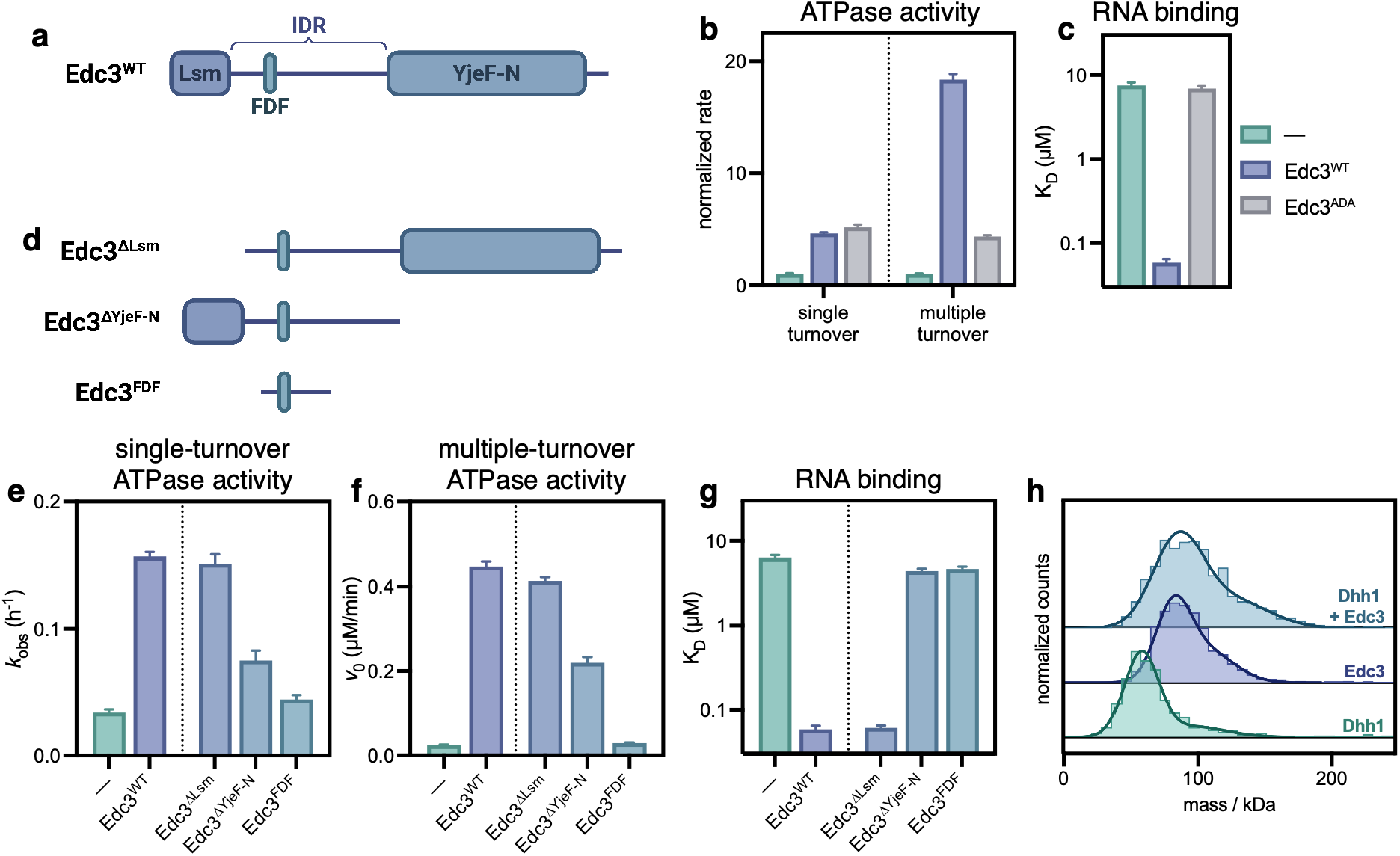
Modulation of Dhh1 activity by Edc3 requires more than the FDF motif. **a.** Domain architecture of Edc3, showing the structured Lsm and YjeF-N domains, the interdomain IDR, and the conditionally structured FDF motif. **b.** Enzymatic activity of Dhh1 under single- and multiple-turnover conditions, in the absence of Edc3 or the presence of Edc3^WT^ and Edc3^ADA^. For each condition, rates are normalized to the rate of Dhh1 alone; both conditions contain saturating U_10_ RNA. **c.** Affinity of Dhh1 for fluorescein-labeled U_20_ RNA in the absence of Edc3 and the presence of Edc3^WT^ and Edc3^ADA^. Fluorescence polarization experiments were performed with 1 mM ADP-BeF_x_. **d.** Domain architecture of Edc3 domain deletion constructs. **e-g.** Effect of Edc3 domain deletion mutants on Dhh1 **(e)** single-turnover and **(f)** multiple-turnover ATPase activity and **(g)** RNA affinity in the presence of 1 mM ADP-BeF_x_. (data for Dhh1 alone and with wild-type Edc3 shown for comparison). **h.** Mass photometry of Dhh1 and Edc3. (Monomeric molecular weights: Dhh1: 55 kDa; Edc3: 49 kDa.) Data in panels (b-c) and (e-g) show fitted rates or affinities from n = 3 technical replicates, with error bars showing standard deviation. See **Tables S3** and **S4** for single-and multiple-turnover ATPase rates shown in panels (b), (e), and (f), **Table S8** for K_D_ values shown in panel (g), and **Table S9** for experimental and theoretical masses for complexes shown in panel (h).

The Edc3 FDF motif is conserved in eukaryotes and interacts with the Dhh1 RecA-C domain to promote translation repression and P-body formation (34, 42, 43, 55–57). A double phenylalanine-to-alanine mutation in the Edc3 FDF motif or mutations in the corresponding Dhh1 RecA-C surface results in significant reduction in co-immunoprecipitation of the *Drosophila* orthologs Edc3 and Me31B (34, 43). Based on these results, we predicted that this double alanine FDF mutation (Edc3^ADA^: F99A/F101A) would abrogate binding to Dhh1 and preclude stimulation of both Dhh1 enzymatic activity and RNA binding. Surprisingly, however, we found that Edc3^ADA^ stimulated the activity of Dhh1 to the same extent as wild-type Edc3 in single-turnover ATPase assays (**Fig. 2b**, **Table S3**). Under multiple-turnover conditions, although Edc3^ADA^ increases Dhh1 activity, it does so to a lesser extent than does Edc3^WT^, which suggests that the FDF motif is involved in promoting the release of ADP and inorganic phosphate (P_i_) (**Fig. 2b**, **Table S4**). Additionally, Edc3^ADA^ does not enhance RNA binding by Dhh1, indicating that the FDF motif is also required for this function (**Fig. 2c**, **Table S8**). Taken together, these results show that mutation of the FDF motif uncouples the effect of Edc3 on the enzymatic activity and the nucleotide-dependent RNA affinity of Dhh1.

The ability of wild-type and Edc3^ADA^ to stimulate Dhh1 activity to comparable levels in single-turnover conditions suggests that both constructs interact with Dhh1 under the conditions of this assay. Indeed, in mass photometry (MP) experiments of binary mixtures of Dhh1 with either Edc3^WT^ or Edc3^ADA^, a single peak corresponding to a Dhh1-Edc3 complex is observed in both cases (**Fig. S2a**, lower panel, **Table S9**). However, the ADA mutation does appear to weaken this interaction: at higher salt concentrations (500 mM NaCl, vs. 150 mM NaCl), a binary mixture of Dhh1 and Edc3^ADA^ reveals two distinct peaks, whereas only a single peak is observed with Edc3^WT^ (**Fig. S2a**, upper panel, **Table S9**). Taken together, these results indicate that the interaction between the Edc3 FDF motif and Dhh1 is important but not essential for complex formation, essential for stimulating RNA binding, and critical for product release following ATP hydrolysis. The fact that mutation of the FDF motif has no effect on the ability of Edc3 to stimulate Dhh1 ATP binding and hydrolysis under single-turnover conditions means other sites of interaction are functionally important, however.

We next asked which other domains of Edc3 are involved in modulating Dhh1 activity. To this end, the following constructs were made: an Lsm domain deletion (Edc3^ΔLsm^), a YjeF-N domain deletion (Edc3^ΔYjeF-N^), and a 30-mer FDF motif-containing peptide (Edc3^FDF^) equivalent to a peptide previously used for co-crystallization with Dhh1 and which contains the conserved FDF, FDK, and Trp motifs (**Fig. 2d**) (33). These were tested for the ability to stimulate both the enzymatic activity and RNA binding of Dhh1 (**Fig. 2e-g**, **Tables S3-4** and **S8**). Edc3^ΔLsm^ performs identically to Edc3^WT^, indicating that the Lsm domain is dispensable for the interaction with Dhh1. Removal of the YjeF-N domain, however, results in significantly attenuated stimulation of Dhh1 ATPase activity in both single- and multiple-turnover conditions, as well as complete loss of the enhancement of RNA binding. This is particularly notable given that the YjeF-N domain has not previously been implicated in interactions with Dhh1. The Edc3^FDF^ peptide stimulated neither Dhh1 enzymatic activity nor RNA binding, consistent with previous studies that have reported no functional modulation of Dhh1 by various FDF motif-containing Edc3 peptides and with the requirement for the YjeF-N domain for RNA binding observed here (**Fig. 2g**) (33, 34). Notably, these results show that the FDF motif is essential but not sufficient for stimulation of Dhh1 RNA binding by Edc3 (Edc3^ADA^ in **Fig. 2c** and Edc3^FDF^ in **Fig. 2g**). These results demonstrate that previously unidentified interaction sites in both the IDR and the YjeF-N domain are essential for the modulation of Dhh1 activity by Edc3.

Although only a single Edc3 protomer can interact with the Dhh1 FDF-binding surface, the importance of the YjeF-N homodimerization domain raised the possibility that Edc3 self-association might be necessary for modulation of Dhh1 activity. To test this, we set out to determine the stoichiometry of this interaction using MP (**Fig. 2h**). Although Dhh1 can self-associate via N- and C-terminal IDRs, at the low concentrations used for MP the protein is monomeric, reflecting the low affinity of these interactions (44). In contrast, Edc3, which has a molecular weight of 49 kDa, appears as a dimer, consistent with previous reports that homodimerization of the YjeF-N domain is a high-affinity, likely low-nanomolar interaction (54). Surprisingly, a mixture of Dhh1 and Edc3 reveals a single peak at around 100 kDa, which is inconsistent with a 1:2 Dhh1:Edc3 heterotrimer but consistent with a heterodimer of Dhh1 and Edc3 (**Fig. 2h**, **Table S9**). These results indicate that the interaction of Edc3 with Dhh1 is mutually exclusive with Edc3 dimerization and that the role of the Edc3 YjeF-N domain in stimulation of Dhh1 activity is separate from its role in self-association. The YjeF-N domain does not appear to be a primary driver of affinity for the Dhh1-Edc3 interaction, however, as Edc3^ΔYjeF-N^ readily associates with Dhh1, as seen by MP (**Fig. S2b**).

Taken together, these results show that the interaction between Dhh1 and Edc3 is more complex than was previously appreciated. While the FDF motif is important for the affinity of this interaction, the Edc3 variant harboring a double alanine mutation in this motif (Edc3^ADA^ in **Fig. 2b**-**c**) is still capable of binding to and stimulating ATPase activity of Dhh1. Additionally, deletion of the YjeF-N domain (Edc3^ΔYjeF-N^ in **Fig. 2e-g**) ablates enhancement of RNA binding and significantly attenuates stimulation of Dhh1 ATPase activity by Edc3. Further truncation of the Edc3 IDR and elimination of the Lsm domain to leave a peptide harboring only previously known interaction motifs (Edc3^FDF^ in **Fig. 2e-g**) fully eliminates stimulation of Dhh1 activity. We conclude that functional modulation of Dhh1 by Edc3 requires multiple domains and regions of Edc3, contrary to the prior understanding of this interaction as comprising only on a minimal set of interfaces surrounding the FDF motif.

### Edc3 modulates Dhh1 activity through multipartite interactions

The results from the Edc3 domain deletions indicate that both the IDR (beyond previously identified interaction motifs) and the YjeF-N domain are necessary for full stimulation of Dhh1 activities (**Fig. 2e-g**). This, along with the fact that YjeF-N-mediated Edc3 homodimerization is incompatible with the interaction with Dhh1, suggests that both the IDR and the YjeF-N domain may interact directly with Dhh1. In the absence of structural information beyond the region immediately adjacent to the FDF motif, we used AlphaFold 3 to predict the Dhh1-Edc3 complex in the presence of U_10_ RNA, ATP, and Mg^2+^ (**Fig. 3a**) (40). In this prediction, Dhh1 adopts an active conformation with the two RecA domains forming composite RNA- and ATP-binding sites, comparable to experimental structures solved for other DBPs (e.g. Vasa, **Fig. S3a**). In the absence of RNA and ATP, AlphaFold 3 predicts that Dhh1 in the Dhh1:Edc3 complex assumes an autoinhibited conformation incompatible with ATP binding, similar to the solved crystal structure of apo Dhh1 (**Fig. S3b**).

**Figure 3.**
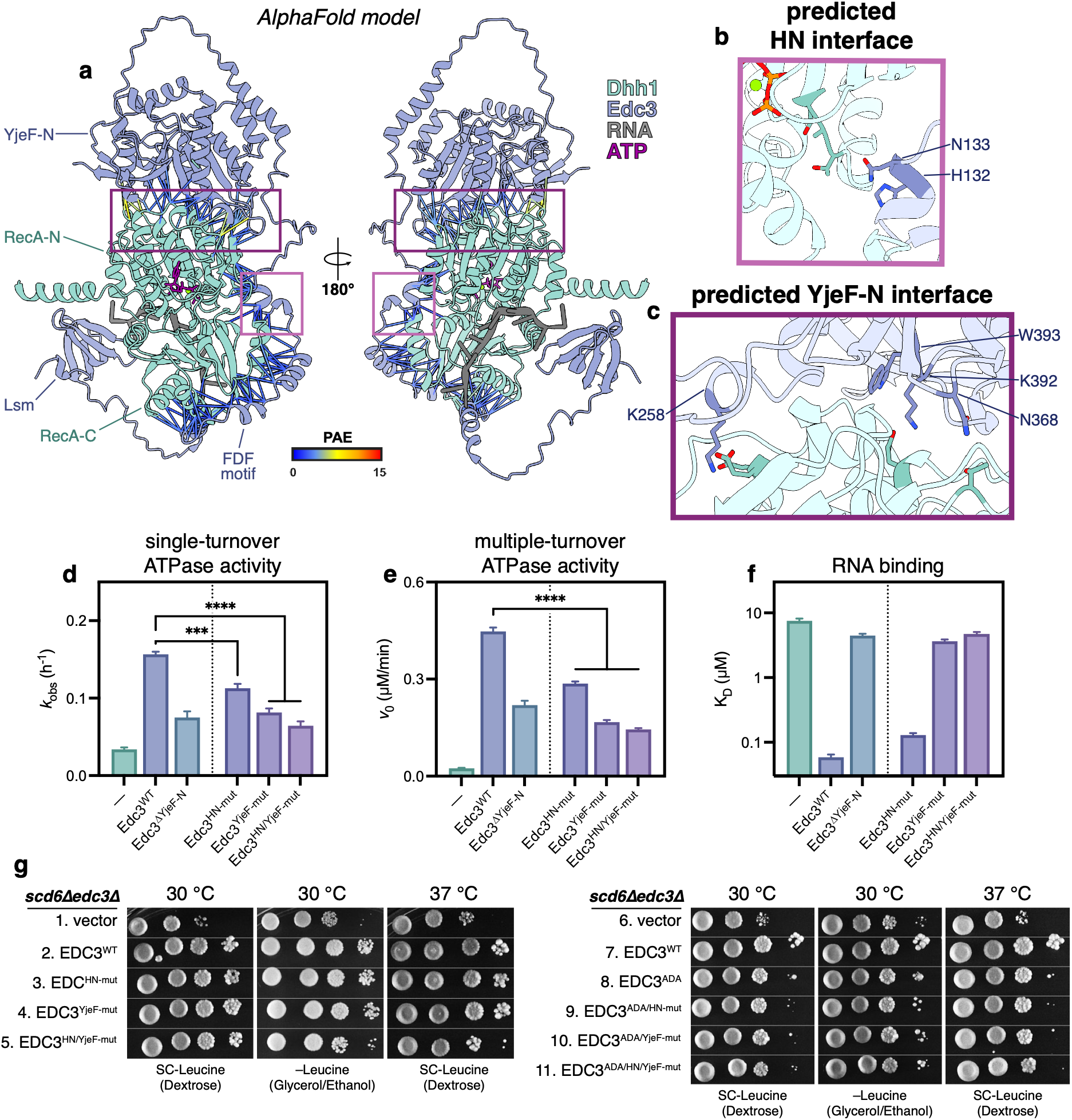
Edc3 interaction with Dhh1 spans multiple domains and motifs. **a.** AlphaFold 3 prediction of interaction between Dhh1 (green) and Edc3 (blue), in the presence of RNA (grey) and ATP (purple). Residues with predicted interprotein distances under 4 Å are shown with connecting lines, colored by Predicted Aligned Error (PAE). Boxed regions show interfaces mutated to make Edc3^HN-mut^ (pink) and Edc3^YjeF-mut^ (purple). **b.** Close-up of HN motif, with residues mutated to make Edc3^HN-mut^ shown in dark blue (H132A/N133A). Dhh1 motif III is highlighted in dark green. **c.** Close-up of YjeF-N interface, with residues mutated to make Edc3^YjeF-mut^ shown in dark blue (K258E/N368A/K392E/W393A). Putative interacting residues on Dhh1 are shown in dark green. **d-f.** Functional effects of mutations at novel predicted interfaces on stimulation by Edc3 of Dhh1 **(d)** single-turnover ATPase activity, **(e)** multiple-turnover ATPase activity, and **(f)** RNA binding. ATPase reactions contain saturating concentrations of U_10_ RNA. Data in panels (d-f) show fitted rates or affinities from n = 3 technical replicates, with error bars showing standard deviation. (Statistical analysis: one-way ANOVA with Dunnett’s multiple comparison test versus Edc3^WT^, *n* = 3) **g.** Cell growth of a transformants of a *scd6ΔedcΔ S. cerevisiae* strain harboring either wild-type or mutant *EDC3* alleles, all tagged with the *myc13* epitope, on single-copy plasmids, grown on plates including or lacking leucine (SC-Leucine and –Leucine, respectively) and incubated at either 30 or 37 °C. Spots represent 10-fold serial dilutions. See **Tables S3** and **S4** for single- and multiple-turnover ATPase rates shown in panels (d) and (e), and **Table S8** for K_D_ values shown in panel (f).

AlphaFold 3 predicts extensive high-confidence interaction interfaces between Dhh1 and Edc3, with low Predicted Aligned Error (PAE) (**Fig. 3a**). Although the conformation of Dhh1 differs depending on whether ATP is included in the prediction, the contacts with Edc3 remain largely the same, likely enabled by the flexibility of the Edc3 IDR (**Fig. S3c-e**). As would be expected, the prediction includes previously identified interaction sites, including the Edc3 FDF, Trp, and FDK motifs (the first two of which are also shared with Scd6/LSM14A) (**Fig. S4**) (31, 33, 34). No high-confidence interactions are predicted between the Edc3 Lsm domain and Dhh1, consistent with the dispensability of this domain for stimulation of Dhh1 (**Fig. 2e-g**). However, in addition to previously reported interactions between Edc3 and the Dhh1 RecA-C domain, the predicted structure also includes novel interactions between the Dhh1 RecA-N domain and both the Edc3 IDR and YjeF-N domains (**Fig. S5a-c**). This led us to hypothesize that multivalent Edc3 interactions spanning both RecA domains may destabilize Dhh1 interdomain interactions, alleviating autoinhibition and facilitating binding of both ATP and RNA (28, 29).

To directly test the functional relevance of the novel predicted interfaces, residues were mutated based on PAE, sequence conservation, and predicted intermolecular interaction (**Fig. S5d-f**): a double mutation of a His-Asn (HN) motif in a short helix in the IDR that is predicted to contact the conserved Dhh1 motif III (Edc3^HN-mut^: H132A/N133A) (**Fig. 3b**); a quadruple mutation at the predicted YjeF-N/RecA-N interface (Edc3^YjeF-mut^: K258E/N368A/K392E/W393A) (**Fig. 3c**); and a combination of the two (Edc3^HN/YjeF-mut^). All three mutants expressed and purified similarly to wild-type Edc3 and interact with Dhh1 with a 1:1 stoichiometry, as seen by MP, indicating that these mutations do not compromise the structure or solubility of Edc3 or its ability to interact with Dhh1, even at low concentrations (**Fig. S6**). However, each mutant displays a compromised ability to modulate the activity of Dhh1. Edc3^HN-mut^ displays moderately decreased stimulation of Dhh1 ATPase activity, while stimulation by Edc3^YjeF-mut^ is attenuated to a similar extent as Edc3^ΔYjeF-N^, indicating that these latter point mutations functionally mimic deletion of the entire domain (**Fig. 3d-e**, **Tables S3-4**). Notably, the effect of these mutations is comparable in both single- and multiple-turnover conditions, unlike mutation of the FDF motif, which exclusively affects stimulation of Dhh1 in multiple-turnover conditions (**Fig. 2b**). The YjeF-N mutations almost completely ablate the effect of Edc3 on Dhh1 binding to RNA, while the HN motif mutations confer no reduction in RNA binding affinity, suggesting distinct stimulatory mechanisms for the interactions with Dhh1 mediated by the HN motif and YjeF-N domain (**Fig. 3f**, **Table S8**). These data are consistent with Edc3 engaging Dhh1 through a multipartite interaction involving the FDF motif, the HN motif, and the YjeF-N domain, with all three interactions contributing to stimulation of Dhh1 ATPase activity, and with the FDF motif and YjeF-N domains being crucial for Edc3-stimulated RNA binding by Dhh1.

To test whether these newly identified interfaces are important for Edc3 function *in vivo*, we performed a comparative growth analysis of *scd6Δedc3Δ* strains of *S. cerevisiae* containing either wild-type or mutant *EDC3* alleles (**Fig. 3g**). Previous work has shown significant functional redundancy between Scd6 and Edc3, such that the double knockout is necessary to observe a growth defect, which can be complemented by either wild-type *SCD6* or *EDC3* on a single-copy plasmid (14). Cell spotting assays were performed in media containing either glucose (SC-Leucine) or nonpreferred carbon sources (–Leucine (Glycerol/Ethanol)) at 30°C or 37°C. Three independent transformants for each *EDC3* allele were examined in parallel; the data shown in **Fig. 3g** are typical for each allele, with additional replicates shown in **Fig. S7**.

The transformants with single-mutant alleles (*EDC3^HN-mut^* and *EDC3^IDR-mut^*) showed growth comparable to the *EDC3^WT^* control in both media and at both 30°C and 37°C (**Fig. 3g**, rows 3, 4 vs. 2). However, the double mutant *EDC3^HN/YjeF-mut^* exhibited slower growth than wild-type and respective single-mutant alleles in both –Leucine media and at 37°C (**Fig. 3g**, row 5 vs. 2-4). These findings suggest that the HN motif and the Dhh1-YjeF-N interface are largely redundant *in vivo*, such that either can be inactivated individually without any detectable effect, but that mutation of both impairs Edc3 function. In contrast, mutation of the Edc3 FDF motif on its own (*EDC3^ADA^*) led to a significant growth defect relative to *EDC3^WT^* (**Fig. 3g**, row 8 vs. 7). However, combining the *HN-mut*, *YjeF-mut*, or *HN/YjeF-mut* mutations with the *ADA* mutation did not produce any additive effects on the observed slow growth phenotype compared to *EDC3^ADA^* alone (**Fig. 3g**, rows 9-11 vs. 8). Importantly, the observed slow-growth phenotype is not attributable to decreased Edc3 expression or stability, as none of the mutants displayed reduced steady-state Edc3 levels, as determined by Western blot analysis (**Fig. S8**). These data are consistent with multipartite interactions between Edc3 and Dhh1 promoting optimal growth in yeast and, moreover, with the interactions observed in *S. pombe* proteins being conserved in *S. cerevisiae*.

### Not1^MIF4G^ and Edc3 stimulate Dhh1 by different mechanisms

Domains with the MIF4G architecture are well-established modulators of DBP activity (59, 38, 60, 61). The MIF4G domain of CNOT1 has previously been shown to bind to DDX6 (the human orthologs of Not1 and Dhh1, respectively), in a manner compatible with simultaneous binding of the CAF1 deadenylase (32). Binding of CNOT1^MIF4G^ was observed to confer substantial rearrangement of DDX6 relative to an apo DDX6 structure, with conserved DBP motifs moving toward canonical active orientations. Based on this, it was previously hypothesized that CNOT1^MIF4G^ stimulates DDX6 activity by relieving autoinhibition and stabilizing a conformation primed for substrate binding (32). Because the DDX6/CNOT1^MIF4G^ structure lacked both RNA and ATP, we sought to test this model by performing kinetic analysis to determine how the MIF4G domain of Not1 (the *S. pombe* ortholog of CNOT1) stimulates Dhh1 activity.

Consistent with previous reports, we find that Not1^MIF4G^ stimulates Dhh1 under both single- and multiple-turnover conditions (**Fig. S9**) (19, 32). In both conditions, Not1^MIF4G^ stimulates Dhh1 to slightly greater extent than does Edc3: 7-fold versus 5-fold in single-turnover conditions, and 51-fold versus 24-fold in multiple turnover conditions (compare **Fig. S1** with **Fig. S9, Tables S3-4**). The fact that greater stimulation is seen under multiple-turnover conditions suggests that Not1^MIF4G^, like Edc3, can also promote a post-hydrolysis step in the Dhh1 catalytic cycle. To determine the mechanism of stimulation of Dhh1 by Not1^MIF4G^, we performed single-turnover ATPase assays over a range of Dhh1 concentrations in the presence of saturating amounts of Not1^MIF4G^ (**Fig. 4a-b**), as was described above for Edc3. In contrast to Edc3, which lowers the K_1/2_ of Dhh1, Not1^MIF4G^ increases the *k*_max_ of Dhh1, while the K_1/2_ remains unchanged from that of Dhh1 alone (**Table S5**). This indicates that Not1^MIF4G^ increases the rate of the chemical step but does not affect the affinity of Dhh1 for ATP. This runs counter to the structure-based hypothesis that binding of CNOT1^MIF4G^ stimulates DDX6 by promoting binding to ATP (32).

**Figure 4.**
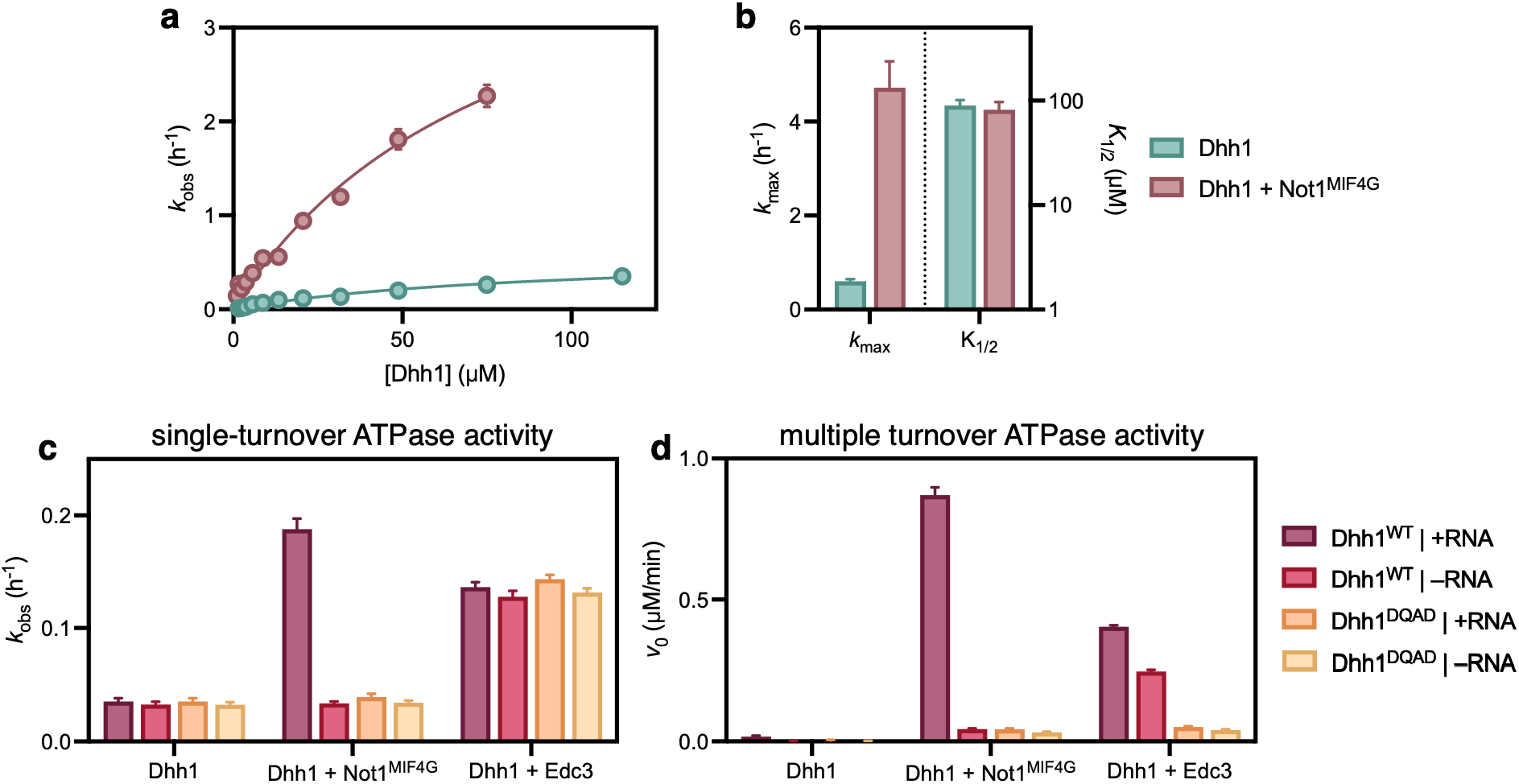
Stimulation of Dhh1 ATPase activity by Not1^MIF4G^ depends on RNA and Walker B glutamate. **a.** Rate of single-turnover Dhh1 ATPase activity as a function of Dhh1 concentration in the absence (green) and presence (red) of saturating amounts of Not1^MIF4G^. All reactions contain saturating amounts of U_10_ RNA. Each point represents fitted rate and standard deviation from n = 3 technical replicates. **b.** Values for *k*_max_ and K_1/2_ of Dhh1 activity in the absence and presence of Not1^MIF4G^, derived from single-turnover kinetic data shown in panel (a), with error bars showing standard deviation. **c-d.** Activity of either Dhh1^WT^ or Dhh1^DQAD^ in both the presence and absence of RNA under **(c)** single-turnover and **(d)** multiple-turnover conditions. Bars show fitted rate with standard deviation from n = 3 technical replicates. See **Table S5** for *k*_max_ and K_1/2_ values shown in panel (b) and **Tables S10** and **S11** for single- and multiple-turnover ATPase rates shown in panels (c) and (d).

DBPs are characterized by nine conserved sequence motifs, with functions related to RNA and ATP binding and ATP hydrolysis. Among these is the Walker B (Asp-Glu-Ala-Asp) motif (also known as Motif II or simply the DEAD motif), which is critical for ATP hydrolysis (25). The glutamate in this motif coordinates the catalytic water molecule, and acts as the general base to promote nucleophilic attack (62–65). We hypothesized that Not1^MIF4G^ increases the maximal rate of Dhh1 by reshaping the active site of Dhh1 to more efficiently hydrolyze ATP. To test this, we next asked whether stimulation of Dhh1 by Not1^MIF4G^ depends on the Walker B motif and the presence of RNA.

Indeed, under single-turnover conditions, removal of RNA or mutation of the Walker B glutamate (Dhh1^DQAD^: D194Q) eliminates the stimulatory effect of Not1^MIF4G^, with ATPase activity reduced to the level of Dhh1 alone (**Fig. 4c**, **Table S10**). This suggests that Not1^MIF4G^ acts cooperatively with RNA and the catalytic glutamate to stabilize the transition state for ATP hydrolysis, thus increasing the *k*_max_. Despite requiring RNA for stimulation of Dhh1, Not1^MIF4G^ neither increases the affinity of Dhh1 for RNA nor induces nucleotide sensitivity for RNA binding (**Fig. S10**). Surprisingly, in the absence of any cofactors, mutation of the conserved Walker B motif glutamate has no effect on the single-turnover ATPase activity of Dhh1 (**Fig. 4c**, **Table S10**). (Although ATP hydrolysis by a DBP harboring a Walker B mutation is highly unusual, it is not without precedent: a DQAD mutant of the *Drosophila* DBP Vasa was previously found be able to hydrolyze ATP (66).) The dispensability of the catalytic glutamate for ATP hydrolysis suggests that, in the absence of Not1^MIF4G^, ATP hydrolysis proceeds through an uncatalyzed, solvent-assisted pathway, rather than through the typical general-base catalyzed mechanism (67, 68). Utilization of a solvent-assisted mechanism could explain the low maximal rate of ATP hydrolysis by Dhh1. Similarly, Dhh1 displays no RNA dependence for ATP hydrolysis in the absence of Not1^MIF4G^, with equivalent rates in both the presence and absence of saturating amounts of U_10_ RNA. In contrast to Not1^MIF4G^, Edc3 stimulates Dhh1^DQAD^ to the same extent as it does wild-type Dhh1, and the effect of Edc3 is not dependent on the presence of RNA (**Fig. 4c**, **Table S10**). This suggests that the binding of Edc3 does not reconfigure the Dhh1 active site to increase the rate of the chemical step, consistent with the fact that Edc3 does not affect the *k*_max_ of Dhh1. Rather, it appears that, when in the presence of Edc3 alone, Dhh1 hydrolyzes ATP through a solvent-assisted pathway that requires neither the Walker B glutamate nor RNA to stabilize the transition state.

Under multiple-turnover conditions, stimulation of Dhh1 by Not1^MIF4G^ is likewise entirely dependent on the Dhh1 Walker B glutamate and RNA (**Fig. 4d**, **Table S11**). With Edc3, however, there is a marked difference between single- and multiple-turnover conditions, with the multiple-turnover activity of Dhh1^DQAD^ much lower than that of wild-type (**Fig. 4d**, **Table S11**). This suggests that the glutamate to glutamine mutation in the Walker B motif inhibits product release, possibly by helping stabilize the hydrolyzed P_i_ in the active site through the loss of charge-charge repulsion and the formation of an additional hydrogen bond, as was suggested previously in the case of Vasa^DQAD^ (66). Under multiple-turnover conditions, the barely measurable activity of Dhh1 alone precludes meaningful analysis of the importance of the Walker B glutamate and the effect of RNA in the absence of cofactors.

We conclude that Not1 and Edc3 act in different steps of the Dhh1 catalytic cycle. Edc3 stimulates Dhh1 ATPase activity by promoting ATP binding, but does not affect the chemical step of catalysis, as seen through the unchanged *k*_max_ (**Fig. 1b**) and by the insensitivity of the rate of ATP hydrolysis to both RNA and mutation of the Walker B glutamate (**Fig. 4c**). In contrast, Not1^MIF4G^ and RNA work together to stimulate the catalytic step of ATP hydrolysis, likely by reconfiguring the Dhh1 active site to allow the Walker B glutamate to coordinate a water molecule, which can then efficiently hydrolyze the ATP γ-phosphate.

### Edc3 and Not1^MIF4G^ cooperatively stimulate Dhh1

Since Edc3 and Not1^MIF4G^ both stimulate Dhh1 ATPase activity, we next asked if these proteins have a combined effect on catalysis. In the presence of both cofactors, single-turnover Dhh1 activity is increased almost 70-fold, as compared to approximately 5- and 7-fold increases in the presence of only either Edc3 or Not1^MIF4G^, respectively (**Fig. S11**, **Table S3**). Under multiple-turnover conditions, Edc3 and Not1^MIF4G^ combine to stimulate Dhh1 activity over 200-fold, again, a much greater increase than was conferred by either cofactor alone (**Fig. 5a-b**, **Table S4**). To better understand the mechanism of this stimulation, we measured the ATPase rate of Dhh1 at a range of concentrations in the presence of saturating amounts of both Edc3 and Not1^MIF4G^ (**Fig. 5c-d**, **Table S5**). Unsurprisingly, given that Edc3 and Not1^MIF4G^ individually modulate either the K_1/2_ or the *k*_max_ of Dhh1, respectively, the two cofactors combine to simultaneously lower the K_1/2_ and increase the *k*_max_. Unexpectedly, however, the magnitude of change for each of these kinetic parameters differs from what was observed with only one cofactor. In the presence of both cofactors, the K_1/2_ of Dhh1 activity is reduced by about 5-fold relative to Dhh1 alone, a smaller effect than that observed in the presence of Edc3 alone, where a 44-fold reduction in K_1/2_ was observed. In the presence of both cofactors, the *k*_max_ increased more than 40-fold relative to Dhh1 alone, which represents a marked increase over the effect of Not1^MIF4G^ in the absence of Edc3, for which an 8-fold increase was observed (**Fig. 5d**, **Table S5**). This suggests that Edc3 enhances the effect of Not1 on Dhh1 activity, possibly by promoting a catalytically productive RNA binding mode which contributes to the Not1-induced RNA- and Walker B-dependent hydrolytic pathway.

**Figure 5.**
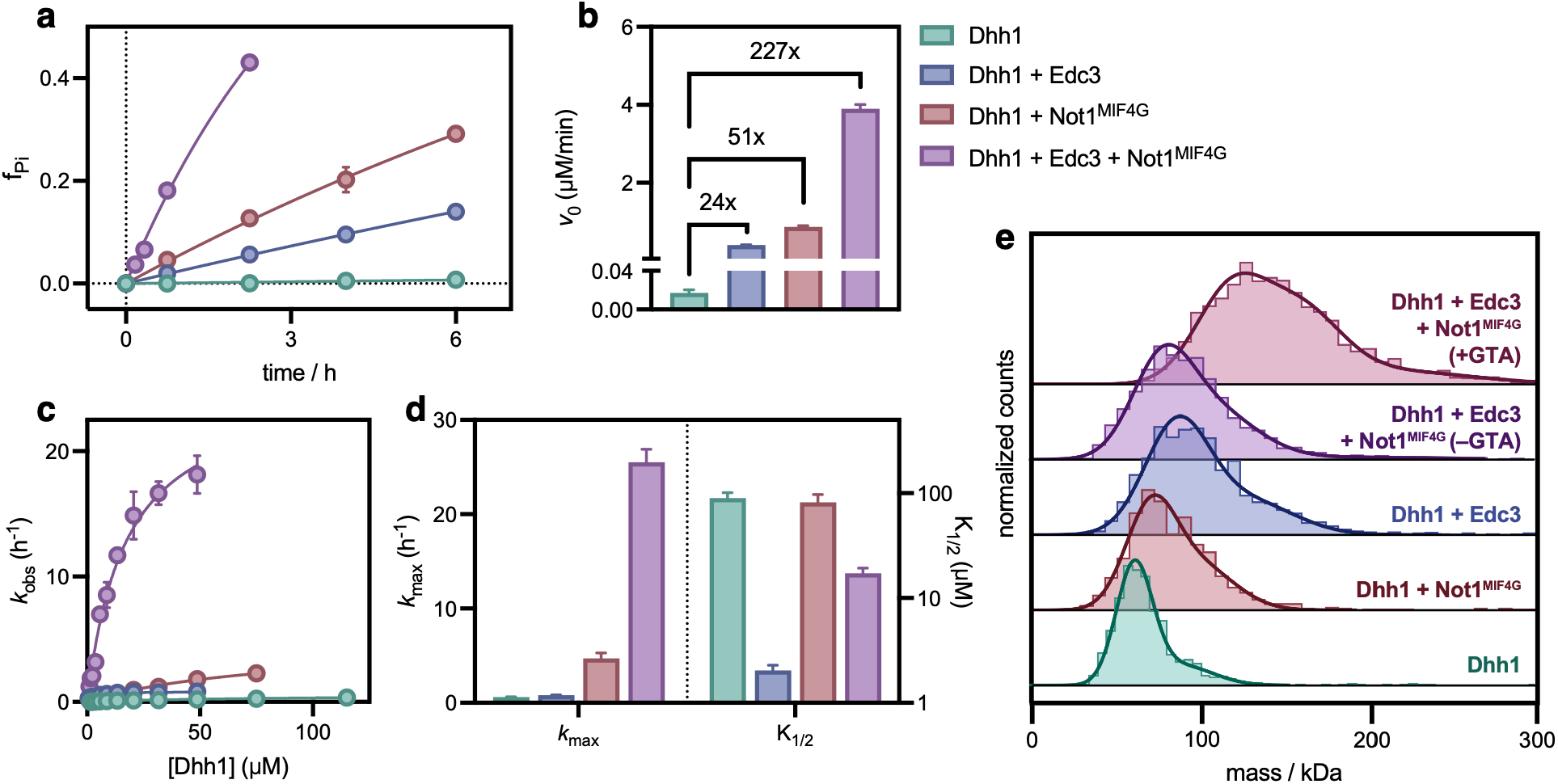
Edc3 and Not1^MIF4G^ cooperatively stimulate Dhh1 activity. **a.** ATPase activity of 5 µM Dhh1 alone (green) and in the presence of 15 µM Edc3 (blue), Not1^MIF4G^ (red), or both Edc3 and Not1^MIF4G^ (pink) under multiple-turnover conditions. Data points show mean and standard deviation of n = 3 technical replicates. **b.** Initial ATPase rates from kinetic data shown in panel (a), with error bars showing standard deviation. **c.** Rate of single-turnover Dhh1 ATPase activity as a function of Dhh1 concentration for Dhh1 alone (green) and in the presence of saturating Edc3 (blue), Not1^MIF4G^ (red), or both Edc3 and Not1^MIF4G^ (pink). All ATPase reactions contain saturating amounts of U_10_ RNA. Each point represents fitted rate and standard deviation from n = 3 technical replicates. **d.** Values for *k*_max_ and K_1/2_ of Dhh1 activity in the absence and presence of Edc3, derived from single-turnover kinetic data shown in panel (c), with error bars showing standard deviation. Data for Dhh1 activity alone and with either Edc3 or Not1^MIF4G^ are repeated from Fig. 1 and Fig. 4 for purposes of comparison. **e.** Mass photometry of equimolar mixture of Dhh1, Edc3, and Not1^MIF4G^ in the presence of U_10_ RNA and ADP-BeF_x_, with and without glutaraldehyde (GTA) crosslinking. See **Table S4** for ATPase rates shown in panel (b), **Table S5** for *k*_max_ and K_1/2_ values shown in panel (d), and **Table S9** for experimental and theoretical masses for complexes shown in panel (e).

Knowing that Not1^MIF4G^ sensitizes Dhh1 to the presence of RNA and configures the catalytic Walker B glutamate, we tested the dependence of Dhh1 activity on these two factors in the presence of both Edc3 and Not1^MIF4G^ (**Fig. S12**, **Tables S10-11**). Removal of RNA or mutation of the Walker B glutamate leads to a reduction in ATPase activity to the level of Dhh1 stimulated by Edc3 alone. This indicates that, in the ternary mixture as in binary mixtures, the effect of Edc3 is totally independent of RNA and the Walker B glutamate, while the effect of Not1^MIF4G^ remains entirely dependent on both. We also tested the affinity of Dhh1 for RNA in the presence of both Edc3 and Not1^MIF4G^ (**Fig. S10b-c**, **Table S7**). In ternary mixtures, as with Edc3 alone, the affinity of Dhh1 for RNA increases by approximately 10- and 100-fold in the presence of ATP and ADP-BeF_x_, respectively, but remains unchanged in the absence of nucleotide or the presence of ADP-AlF_4_ and ADP, indicating that the presence of Not1^MIF4G^ does not meaningfully change the stimulatory effect of Edc3 on Dhh1 binding to RNA.

The fact that the joint effect of Edc3 and Not1^MIF4G^ is not purely additive suggests active synergy between the two cofactors in stimulating the catalytic rate, despite slight antagonistic effects on ATP binding affinity. This result is surprising, as it was previously reported that CNOT1^MIF4G^ and an FDF motif-containing Edc3 peptide do not form a stable heterotrimeric complex with DDX6 (69). As we have already shown that an FDF motif-containing Edc3 peptide does not capture the full extent of the interactions between Edc3 and Dhh1, we used AlphaFold 3 to predict the structure of the heterotrimeric Dhh1/Edc3/Not1^MIF4G^ complex in the presence of U_10_ RNA, ATP, and Mg^2+^ (**Fig. S13**). High-confidence interactions are predicted between Dhh1 and both Edc3 and Not1^MF4G^, with essentially no high-confidence interactions predicted between Edc3 and Not1^MIF4G^ (**Fig. S13c**). The predicted interfaces of Edc3 and Not1^MIF4G^ with Dhh1 are highly similar to those found in the predicted Dhh1/Edc3 dimer and the experimental DDX6/CNOT1^MIF4G^ structures, respectively (**Fig. S13d-i**). This prediction suggests that, even with full-length Edc3, the binding sites on Dhh1 for Edc3 and Not1^MIF4G^ are spatially distinct.

We hypothesized that, although an Edc3^FDF^ peptide does not form a stable heterotrimeric complex with DDX6 and CNOT1^MIF4G^, full-length Edc3 might be able to form a stable complex with Dhh1 and Not1^MIF4G^ due to the additional points of contact relative to the peptide. Using MP to assess complex formation in the presence of U_10_ RNA and ADP-BeF_x_, we found that, while both Edc3 (as previously shown) and Not1^MIF4G^ form stable dimers with Dhh1, no stable heterotrimeric complex could be observed (**Fig. 5e**, **Table S9**). Instead, in a mixture of all three proteins, only a peak around 100 kDa was observed, likely representing the Dhh1/Edc3 heterodimer, with Not1^MIF4G^ unbound (at 30 kDa, Not1^MIF4G^ is below the detection limit of this technique and so cannot directly be observed). We reasoned that, although Dhh1, Edc3, and Not1^MIF4G^ do not form a stable complex, the co-modulation of Dhh1 activity by Edc3 and Not1^MIF4G^ might instead be due to the formation of a transient, functional heterotrimeric complex. To test this, we performed the same MP experiment, with the addition of glutaraldehyde to crosslink and trap any transient interactions. Upon glutaraldehyde crosslinking, we observe a shift to a peak around 130 kDa, consistent with the expected mass of the heterotrimeric complex (134 kDa), suggesting that Edc3 and Not1^MIF4G^ transiently co-engage Dhh1, likely leading to the strong synergistic stimulation of ATPase activity (**Fig. 5e**, **Table S9**). To ensure that the observed mass shift is not due to non-specific interactions between any of the proteins in the ternary mixture, we performed crosslinking experiments for all binary combinations of Dhh1, Edc3, and Not1^MIF4G^ (**Fig. S14**). A mass shift upon glutaraldehyde addition was not observed for any of the three binary mixtures (panels a-c), indicating that this is specific to the ternary mixture (panel d) and that the observed increase in mass represents a Dhh1/Edc3/Not1^MIF4G^ heterotrimeric complex.

## DISCUSSION

Dhh1 is a highly abundant DBP that plays a key role in various RNA-related processes, including translational repression, P-body formation, and RNA decay. Despite well-established functions *in vivo*, the molecular mechanism of Dhh1 is poorly understood. Here, we show that the 5’-3’ decay factors and previously known Dhh1 interactors Edc3 and Not1 combine to modulate both the RNA binding and ATPase functions of Dhh1 in a manner that brings the activity of the enzyme more in line with canonical behavior of the DBP family. Edc3 both increases the enzymatic activity of Dhh1 by enhancing binding of ATP and enhances the affinity of Dhh1 for RNA in a nucleotide-dependent manner. Surprisingly, we find that this modulation of activity is the product of extensive, domain-spanning interactions, whereas prior studies identified only a minimal set of small interaction motifs (the FDF, FDK, and Trp motifs) as contributing to the Dhh1-Edc3 interaction (33, 34, 43, 57). In contrast to Edc3, the MIF4G domain of Not1 increases the activity of Dhh1 by increasing the maximal rate of the enzyme in a manner fully dependent on RNA and the Walker B motif, suggesting that Not1 remodels the active site in conjunction with RNA to engage the conserved Walker B glutamate to facilitate ATP hydrolysis. Moreover, Edc3 and Not1^MIF4G^ combine to synergistically stimulate Dhh1 catalytic activity, apparently through formation of a short-lived functional heterotrimeric complex.

Here, we will discuss the following: first, we will propose a model of cofactor-regulated Dhh1 enzymatic activity; second, we will compare our results with general models of DBP activity; third, we will examine how our observations expand what is known about the Dhh1-Edc3 interaction both structurally and functionally; fourth, we will discuss the implications of our results for the interactions of Dhh1 with other proteins; and, finally, we will speculate on the role of Dhh1 activity in 5’-3’ RNA decay.

### Model of Edc3- and Not1-mediated Dhh1 activity

Based on our data, we propose the following model for the cofactor-regulated enzymatic cycle of Dhh1 activity (**Fig. 6, i-viii**). Alone, Dhh1 has a low affinity for both ATP and RNA (**Fig. 1d** and **g**). (**i**) Binding of Edc3, spanning both RecA domains, alleviates inhibitory intradomain interactions in Dhh1 and (**ii**) primes the enzyme for binding ATP with much higher affinity (**Fig. 1d**). (**iii**) In the Edc3- and ATP-bound state, the affinity of Dhh1 for RNA increases substantially (**Fig. 1g**); this Dhh1/Edc3/RNA complex, while stable, still has a low maximal rate of ATP hydrolysis, likely due to utilization of an inefficient solvent-assisted mechanism (**Fig. 1b**). (**iv**) Binding of Not1 via the MIF4G domain allosterically reconfigures the Dhh1 active site (in combination with the already-bound RNA) to orient the Walker B glutamate to coordinate a nucleophilic water molecule, thus increasing the maximal rate of the enzyme (**Fig. 4c-d** and **Fig. 5d**), (**v**) leading to ATP hydrolysis. (**vi**) The Dhh1/Edc3/Not1 complex is unstable, with Not1 rapidly dissociating (**Fig. 5e**). (**vii**) Following ATP hydrolysis, the Edc3-, ADP- and P_i_-bound Dhh1 has a reduced affinity for RNA, leading to RNA release (**Fig. 1g**). (**viii**) Edc3 facilitates the release of ADP and P_i_, which otherwise form an inhibitory and likely high-affinity complex with Dhh1 (**Fig. S1**). Following the release of the ATP hydrolysis products, the Dhh1/Edc3 complex is again primed to undergo the catalytic cycle again.

**Figure 6.**
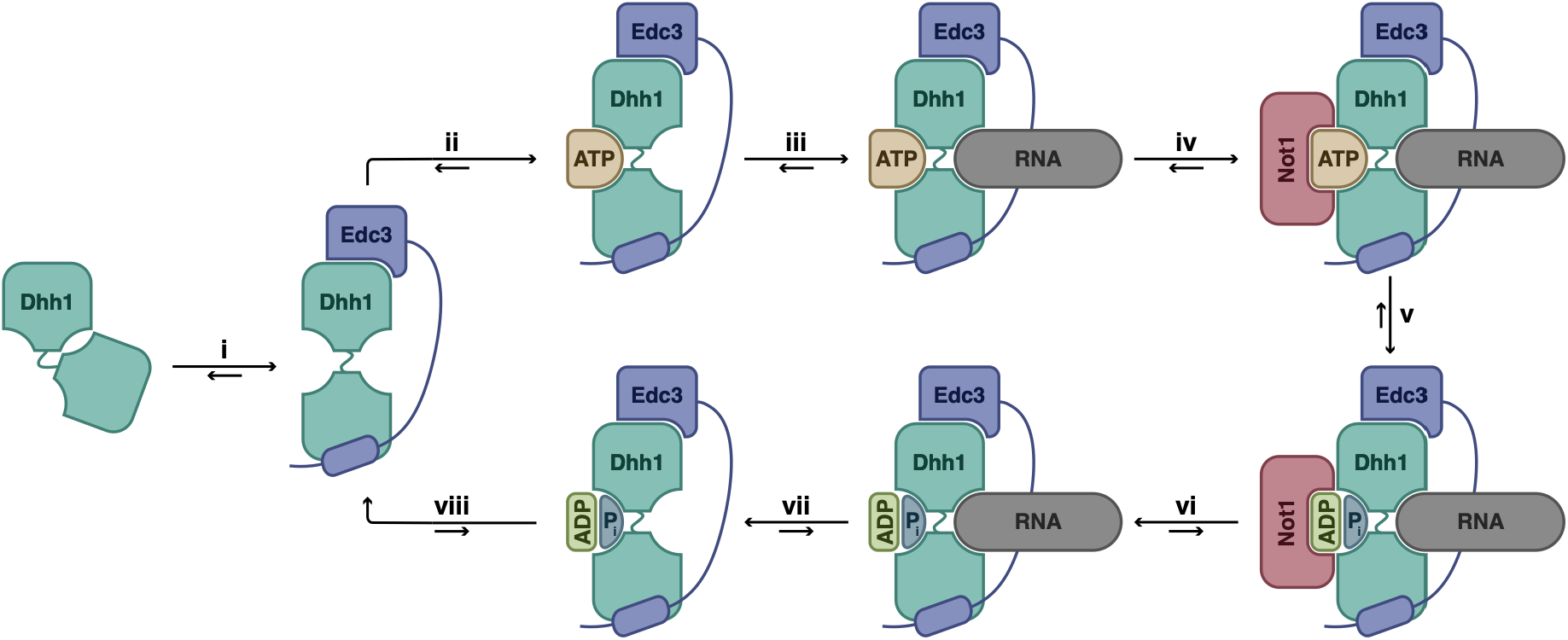
Edc3- and Not1-mediated catalytic cycle of Dhh1. Proposed model showing how Edc3 and Not1 regulate Dhh1 activity, showing *(i)* Edc3 binding, *(ii)* ATP binding, *(iii)* RNA binding, *(vi)* Not1 binding, *(v)* ATP hydrolysis, *(vi)* Not1 unbinding, *(vii)* RNA release, and *(viii)* release of ADP and P_i_.

### Comparison between Dhh1 and other DBPs

Prior studies have shown that proteins with HEAT repeat-containing MIF4G domains elicit varied effects on DBP function, with eIF4G and Gle1 stimulating the activity of eIF4A and Dbp5, respectively, whereas CWC22 locks eIF4AIII into an inactive state (36, 59, 60). Like eIF4G and Gle1, the MIF4G domain of Not1 has been shown to stimulate the ATPase activity of Dhh1 (19, 32). For eIF4G, stimulation of eIF4A is achieved through both increasing the affinity of eIF4A for ATP and increasing the *k*_cat_ of ATP hydrolysis (70, 71). In contrast, we show here that Not1^MIF4G^ only affects the chemical step of ATP hydrolysis, without changing substrate affinity (**Fig. 4b**). Instead, there is an apparent division of labor between Edc3 and Not1 for regulation of the Dhh1 catalytic cycle: Edc3 likely opens the composite ATP binding site in Dhh1 to promote nucleotide and attendant RNA binding, while Not1 reconfigures the active site to accelerate the chemical step. These results underscore that, despite the conserved structural features of both DBPs and MIF4G domains, the functional and mechanistic effects of DBP-MIF4G interactions are remarkably diverse.

Two particularly surprising features of Dhh1 shown here are the dispensability of the Walker B glutamate and the lack of stimulatory effect of RNA for catalyzing ATP hydrolysis in the absence of Not1^MIF4G^ (**Fig. 4c-d**). The activity of DBPs is, almost as a rule, stimulated by RNA by as much as several orders of magnitude (27); likewise, the Walker B glutamate is universally conserved and critical for function not only within the DBP family, but in the greater SF1/SF2 helicase superfamilies (72).

Since mutation of the conserved Walker B motif glutamate has no effect on the single-turnover ATPase activity of Dhh1, we conclude that, in the absence of Not1^MIF4G^, ATP hydrolysis proceeds through a pathway independent of general base catalysis regardless of whether Edc3 is present (**Fig. 4c**). Under multiple turnover conditions, when ATP is in excess of enzyme, however, the DQAD mutant is significantly less active than wild-type (in the presence of Edc3; alone, the multiple-turnover rate of Dhh1 is too low to be able to observe a decrease in activity) (**Fig. 4d**). This suggests that this mutation prevents the release of ADP and P_i_, even though it does not prevent ATP hydrolysis. This phenomenon was previously observed with the *Drosophila* DBP Vasa (DDX4 in humans), for which this same DQAD Walker B mutation did not affect ATP hydrolysis under single-turnover conditions but significantly reduced activity under multiple-turnover conditions (66). Consistent with this mutant being competent for ATP hydrolysis but displaying impaired product release, crystallization of Vasa^DQAD^ in the presence of ATP revealed trapped hydrolysis products ADP and P_i_ in the active site (66). Notably, Vasa, like Dhh1, has an unusually low ATP turnover rate, suggesting that it may also hydrolyze ATP through a general base-independent, solvent-assisted pathway in the absence of stimulating cofactors (68, 73, 74).

The lack of dependence on the Walker B glutamate for single-turnover hydrolysis and the lack of stimulation by RNA for both Dhh1 and Vasa may be linked. In other DBPs, both RNA and the catalytic glutamate are critical for ATPase *k*_cat_, with removal of the former or mutation of the latter resulting in unchanged ATP affinity but decreased maximal hydrolysis rates (45, 49, 75). We have shown here that for Dhh1, the addition of Not1^MIF4G^ increases the *k*_max_ while also sensitizing Dhh1 to the presence of RNA and to mutation of the Walker B glutamate. The absence of any of Not1^MIF4G^, RNA, or the Walker B glutamate is sufficient to return Dhh1 to its low basal activity, with the absence of multiple elements causing no additional reduction in rate. This interdependent cooperativity suggests that Not1^MIF4G^ and RNA increase the maximal rate of Dhh1 by reorienting the catalytic glutamate, thus stabilizing an energetically favorable transition state for ATP hydrolysis. Allosteric modulation of the Walker B motif to properly configure the catalytic glutamate may be a generalized feature of stimulation of DBP ATPase activity by RNA. Comparison of crystal structures of the human DBP DDX19 with the ATP analog AMP-PNP in either the presence or the absence of RNA shows that, in the presence of RNA, DDX19 adopts a closed conformation with the Walker B glutamate oriented to coordinate the nucleophilic water, while in the absence of RNA, the enzyme adopts an open conformation with the Walker B glutamate oriented away from the bound nucleotide (**Fig. S15**).

### New insights into the Dhh1-Edc3 interaction

Prior studies have established that the FDF motif is important for the interaction between Edc3 and Dhh1, with mutations at this interface being used to disrupt binding *in vivo* in order to probe the functional relevance of the interaction (30, 33, 34, 43, 76, 77). Unexpectedly, we found that, although the FDF motif might be necessary for the interaction between Dhh1 and Edc3 *in vivo*, it is neither necessary for the interaction *in vitro* nor sufficient for functional modulation of Dhh1 by Edc3 (**Fig. 2b-c** and **e-g**). We show that, in addition to the previously identified FDF, FDK, and Trp motifs, the YjeF-N domain and the HN motif are functionally important for the Dhh1-Edc3 interaction. Interestingly, mutation of these different interaction surfaces lead to different functional outcomes: the FDF motif and the YjeF-N domain are necessary for nucleotide-dependent RNA binding; the HN motif and the YjeF-N domain are important for single-turnover activity; and all three are involved in promoting the release of ADP and P_i_. Intriguingly, the YjeF-N domain is critical for the stimulation of Dhh1 ATPase activity and RNA binding, despite the YjeF-N being predicted to contact the Dhh1 RecA-N domain, distal from both the ATP- and RNA-binding pockets. It is possible that this distal interaction serves to help alleviate previously observed autoinhibitory interactions between the Dhh1 RecA domains, although further experimentation is required to directly test that hypothesis (28, 29). Importantly, mutation of both the HN and YjeF-N interfaces led to growth defects in a *scd6Δedc3Δ* strain of *S. cerevisiae*, indicating the functional relevance of these novel Edc3-Dhh1 interfaces *in vivo* and the conservation of these interactions between *S. pombe* and *S. cerevisiae* (**Fig. 3g**).

Mutation of any single interaction surface does not prevent Dhh1-Edc3 heterodimerization, even at double-digit nanomolar concentrations, suggesting that no individual motif is responsible for the high-affinity interaction (**Fig. S2a** and **Fig. S6b**). That each of these interaction motifs have different functional effects on Dhh1 activity suggests that the Dhh1-Edc3 interaction may be quite dynamic, with different domains and motifs interacting in turn at different points of the Dhh1 catalytic cycle. However, the FDF, FDK, and Trp motifs likely contribute significantly to the affinity of this interaction, with a reported nanomolar affinity for an 81-residue *S. cerevisiae* Edc3 peptide including these motifs (33). It is possible that these motifs serve to anchor the Dhh1-Edc3 interaction, with the HN motif and YjeF-N domain interacting subsequently to modulate Dhh1 activity. This model is consistent with the strong slow growth phenotype seen for *EDC3^ADA^* and the fact that additional mutations at the HN motif or YjeF-N interface produced no additive effects to the *EDC3^ADA^* slow growth (**Fig. 3g**).

We have shown here that the binding of Edc3 to Dhh1 has significant functional implications for Dhh1 function, with RNA binding, ATP binding, ATPase rate, and ADP release all affected. The observation that Edc3 enhances the affinity of Dhh1 for RNA is somewhat surprising, as a previous study reported that an FDF motif-containing Edc3 peptide disrupts binding of Dhh1 to RNA in both the absence of nucleotide and in the presence of the ATP analog AMP-PNP (33). This discrepancy can be explained in part by the observation that full-length Edc3 has a minimal effect on the affinity of Dhh1 for RNA in the presence of AMP-PNP (**Fig. S16**). This likely reflects the fact that AMP-PNP can adopt various active-site conformations, some of which mimic the post-hydrolysis, rather than pre-hydrolysis, state (78). Moreover, we have shown here that an FDF motif-containing Edc3 peptide alone is insufficient to stimulate RNA binding by Dhh1 even in the presence of ADP-BeF_x_ (**Fig. 2g**). As such, we conclude that these results are not contradictory but rather are likely simply due to differences in protein constructs, nucleotide analogs, and experimental methods.

The *in vivo* relevance of the canonical DBP activities of Dhh1 has previously been established by mutating conserved DBP motifs of known function. Mutations in motifs necessary for RNA and ATP binding leads to defects in Dhh1-dependent RNA decay, cell growth sensitivity to temperature and stress, as well as deficiency in P-body formation (13, 19, 28, 29). Similarly, mutations to residues in Dhh1 important for ATP hydrolysis negatively affect RNA decay and cell growth, lead to deficiencies in Dhh1-related translational repression, and also prevent the dissolution of P-bodies (19, 28, 29, 32, 76, 79). Here we show that Edc3 affects these same critical biochemical functions (ATP and RNA binding, ATP hydrolysis) *in vitro*, suggesting that Edc3 may contribute to some of these *in vivo* processes by directly modulating Dhh1 activity. Edc3 has been shown to act (redundantly with Scd6) to recruit Dhh1 to the decapping complex, likely to help target Dcp2 to substrates, a process that may require the RNA binding and ATP binding activities of Dhh1 (14, 35).

### Implications for other Dhh1 interactors

In addition to Edc3, Dhh1 is known to interact with other proteins via FDF (or pseudo-FDF) motifs, including the RNA decay factors Pat1 and Scd6 (Trailer Hitch/Tral in *Drosophila*, LSM14A in humans), the translational repressors 4E-T and GIGYF, and the large ribosomal subunit RPL22 (31, 33, 34, 69, 76, 80). In the 5’-3’ decay pathway, the mutually exclusive nature of the interactions with Edc3, Scd6, and Pat1 has led to the proposal that these activators bind competitively to Dhh1 to promote the degradation of specific transcripts (35, 81).

Given that Edc3 actively modulates Dhh1 activity, it is possible that these other FDF motif-containing proteins also affect Dhh1 function beyond simple binding and recruitment. Other interactors may differ in their effects on Dhh1 function since, as shown here, interactions beyond the Edc3 FDF motif are essential for modulation of Dhh1 activity. Scd6 has recently been shown to exhibit significant functional redundancy with Edc3, raising the possibility that it also binds Dhh1 across multiple domains to modulate RNA binding and ATPase activity (14). Although Scd6 is predicted to be largely unstructured and lacks a domain analogous to the Edc3 YjeF-N domain, a high-confidence interaction is predicted between Scd6 and the Dhh1 motif III, analogous to the Dhh1 interaction involving the Edc3 HN motif (**Fig. S17**). Similarly, other proteins that interact with Dhh1 via an FDF motif may contain additional regions that interact with Dhh1 to tune RNA binding, ATP binding, or ATPase activity.

Additionally, these results suggest new interaction modes and novel interaction surfaces for Dhh1 that could play diverse roles in its various biological functions. Previously, direct interactions have centered around relatively few sites on Dhh1, namely the surfaces that have been previously shown to interact with the FDF, FDK, and Trp motifs, as well as the Not1^MIF4G^ binding site (see **Fig. S4** and **Fig. S11**, respectively). Based on immunoprecipitation-mass spectrometry data, the human Dhh1 ortholog DDX6 was reported to make at least 29 reported direct interactions, excluding ribosomal proteins (the actual total is certainly higher, as this experiment missed some known interactors, including CNOT1 and GIGYF1) (82). Of these 29 identified proteins, only five have an identifiable FDF motif. That Edc3 makes important contacts with other surfaces on Dhh1, including the interdomain region close to motif III and the RecA-N domain, suggests that other direct Dhh1/DDX6 interactors may make use of these or similar surfaces.

### Role of Dhh1 in bridging deadenylation and decapping in 5’-3’ RNA decay

Dhh1 is known to act between the deadenylation and decapping steps of the 5’-3’ mRNA decay pathway (7). The binary interactions between Dhh1 and both Edc3 (as well as additional FDF motif-containing proteins Pat1 and Scd6) and Not1 have led to the hypothesis that Dhh1 serves to physically link the Ccr4-Not deadenylase complex to the decapping complex (69, 81). The fact that Dhh1 activity is tuned by Not1 and Edc3 suggests that, beyond simply binding and releasing various decay factors, Dhh1 activity may be directly involved in bridging 3’ deadenylation and 5’ decapping. Consistent with this, mutations in Dhh1 that disrupt either RNA or ATP binding or ATP hydrolysis lead to stabilization of Dhh1-sensitive transcripts (28, 29, 76).

Based on the data presented here, it is possible to hypothesize mechanisms by which Dhh1 might bridge deadenylation and decapping. For example, Edc3- and ATP-bound Dhh1 could recruit the decapping complex (which binds to the Edc3 Lsm domain) to transcripts in a sequence independent manner via its high affinity interaction with RNA. Interaction with Not1 could then trigger the hydrolysis of ATP, leading to release of RNA by Dhh1, thus liberating the decapping complex to bind to and subsequently remove the 5’ cap. Consistent with this, previous work has shown that Dhh1-sensitive transcripts are stabilized by the DQAD mutation, but that Dhh1^DQAD^ shows no defects in promoting decay when artificially targeted to a transcript (13, 76). This suggests that the function of Dhh1 in 5’-3’ decay requires ATPase activity only insofar as it affects RNA binding, likely through the nucleotide-coupled changes in RNA affinity shown here. Further work, both *in vitro* and *in vivo*, is needed to fully understand the role of Dhh1 in bridging deadenylation and decapping, as well as how modulation of Dhh1 activity by Not1 and Edc3 contribute to this process. Among the outstanding questions is the effect on Dhh1 function of additional deadenylation and decapping factors and complexes, including full-length Not1, the complete Ccr4-Not complex, decapping complexes including the Dcp1/Dcp2 decapping enzyme, and other factors such as Scd6 and the Pat1/Lsm1-7 complex. Additionally, the studies here used purified proteins and short model RNAs; further investigation is required to understand how the effects shown here translate to the lower concentrations, diverse RNA substrates, and complex macromolecular environment of the cytoplasm to fully understand the role of Dhh1 activity in 5’-3’ mRNA decay.

## ACKNOWLEDGMENTS

We thank Estelle Ronayne for her insightful comments on the manuscript. We thank the lab of Lori Passmore (University of Cambridge) for providing the Not1^MIF4G^ plasmid.

## AUTHOR CONTRIBUTIONS

Conceptualization: G.A.B and J.D.G.; data curation: G.A.B. and J.D.G.; formal analysis: G.A.B. and R.K.; funding acquisition: G.A.B., A.G.H., and J.D.G.; investigation: G.A.B. and R.K.; methodology: G.A.B.; project administration: J.D.G.; resources: A.G.H. and J.D.G.; supervision: A.G.H. and J.D.G.; validation: G.A.B. and R.K.; visualization: G.A.B.; writing – original draft: G.A.B.; writing – review & editing: G.A.B., A.G.H., J.D.G.

## SUPPLEMENTARY DATA

Supplementary data is available in the accompanying file.

## DATA AVAILABILITY

The data underlying this article are publicly available at https://doi.org/10.5281/zenodo.19666725.

