## Supplementary Information for "Deadenylation and Decapping Factors Cooperatively Stimulate Biochemical Activities of DEAD-Box ATPase Dhh1"

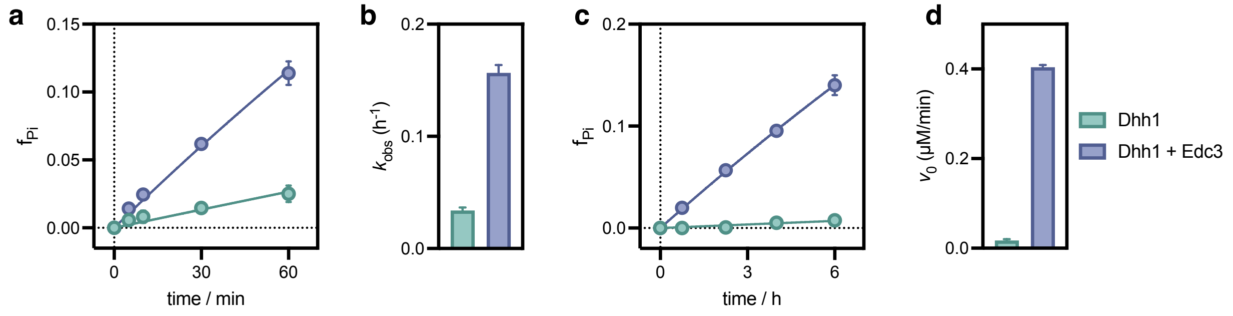


**Figure S1. Edc3 stimulates Dhh1 ATPase under single- and multiple-turnover conditions**
ATPase activity of 5 $\mu$M Dhh1 in the absence (green) and presence (blue) of 15 $\mu$M Edc3 under **(a-b)** single-turnover and **(c-d)** multiple-turnover conditions. Total ATP concentration is 1 nM and 1 mM, respectively, and both conditions contain saturating amounts of $U_{10}$ RNA. Data points in panels (a) and (c) show mean and standard deviation from n = 3 technical replicates; error bars in panels (b) and (d) show standard deviation of fitted rates. See **Tables S3** and **S4** for single- and multiple-turnover ATPase rates shown in panels (b) and (d).

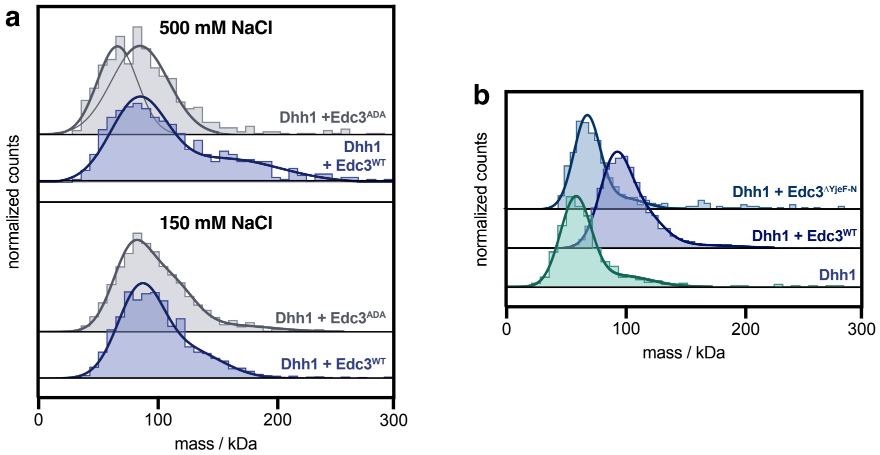


**Figure S2. Edc3$^{ADA}$ and Edc3$^{YjeF-N}$ form complexes with Dhh1**

**a.** Mass photometry of binary mixtures of Dhh1 and either Edc3$^{WT}$ (blue) or Edc3$^{ADA}$ (grey), under standard salt (150 mM NaCl) and high salt (500 mM NaCl) conditions. **b.** Mass photometry of binary mixture of a binary mixture of Dhh1 and Edc3$^{\Delta YjeF-N}$ (light blue); Dhh1 alone (green) and Dhh1 with Edc3$^{WT}$ (dark blue) are included for reference. (Monomeric molecular weights: Dhh1: 55 kDa; Edc3$^{WT}$ and Edc3$^{ADA}$: 49 kDa; Edc3$^{\Delta YjeF-N}$: 21 kDa.) See **Table S9** for experimental and theoretical masses for complexes shown in both panels.

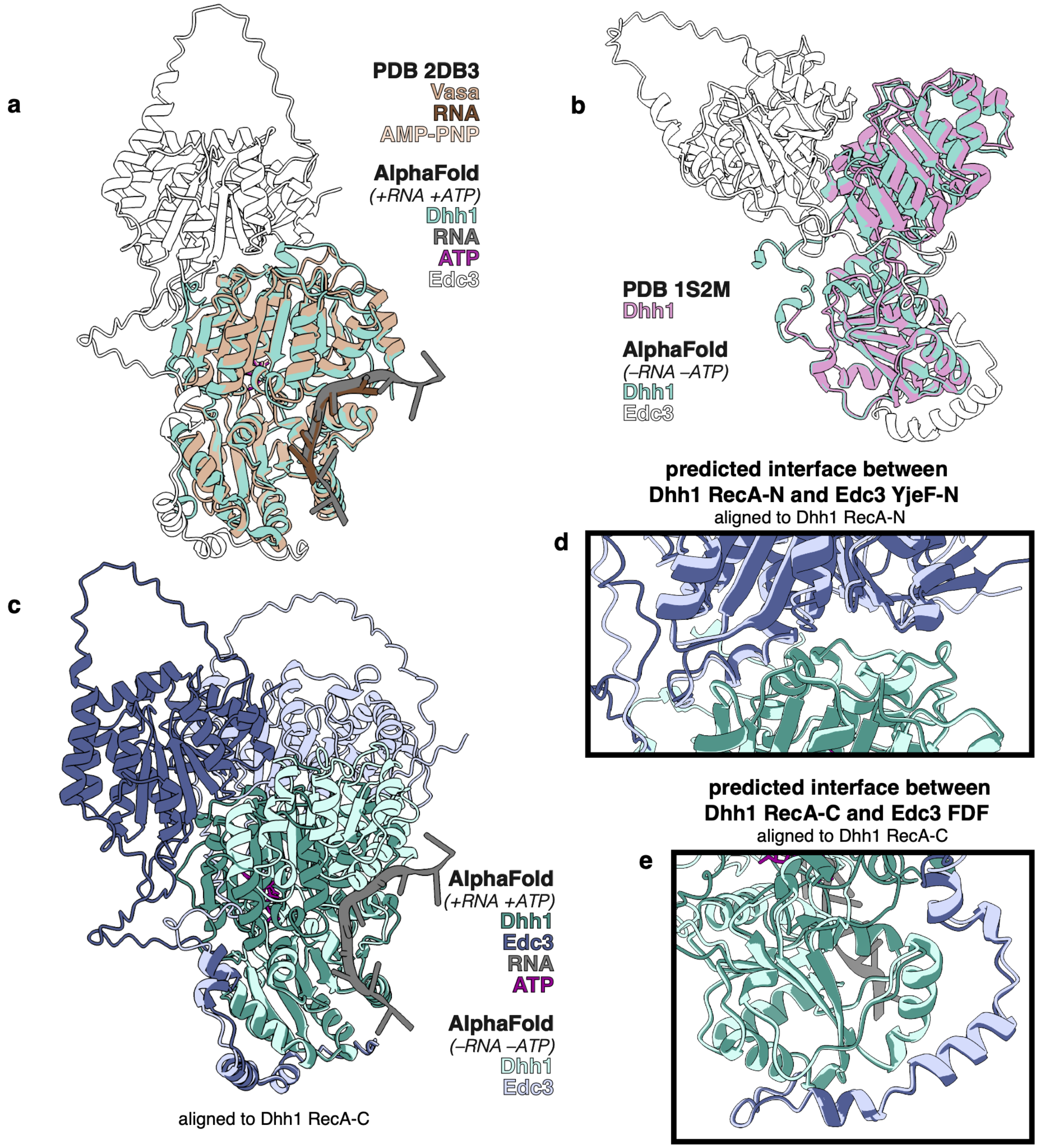


**Figure S3. Dhh1-Edc3 structural predictions in the presence and absence of RNA and ATP**
**a.** Predicted Dhh1-Edc3 dimer in the presence of RNA and ATP, compared to experimental structure of *Drosophila* DBP Vasa in active state (PDB: 2DB3). **b.** Predicted Dhh1-Edc3 dimer in the absence of RNA and ATP, compared to experimental structure of Dhh1 in inactive state (PDB: 1S2M). **c.** Prediction of Dhh1/Edc3 heterodimer in the presence (dark colors) and absence (light colors) of RNA and ATP. **d-e.** Isolated interaction interfaces are similar regardless the presence of RNA and ATP, including the interaction between **(d)** Edc3 YjeF-N domain and Dhh1 RecA-N and **(e)** Edc3 FDF region and Dhh1 RecA-C. Coloring is the same as panel (c). For all predicted structures, the Dhh1 N- and C-terminal IDRs and the Edc3 Lsm domain are hidden for clarity.

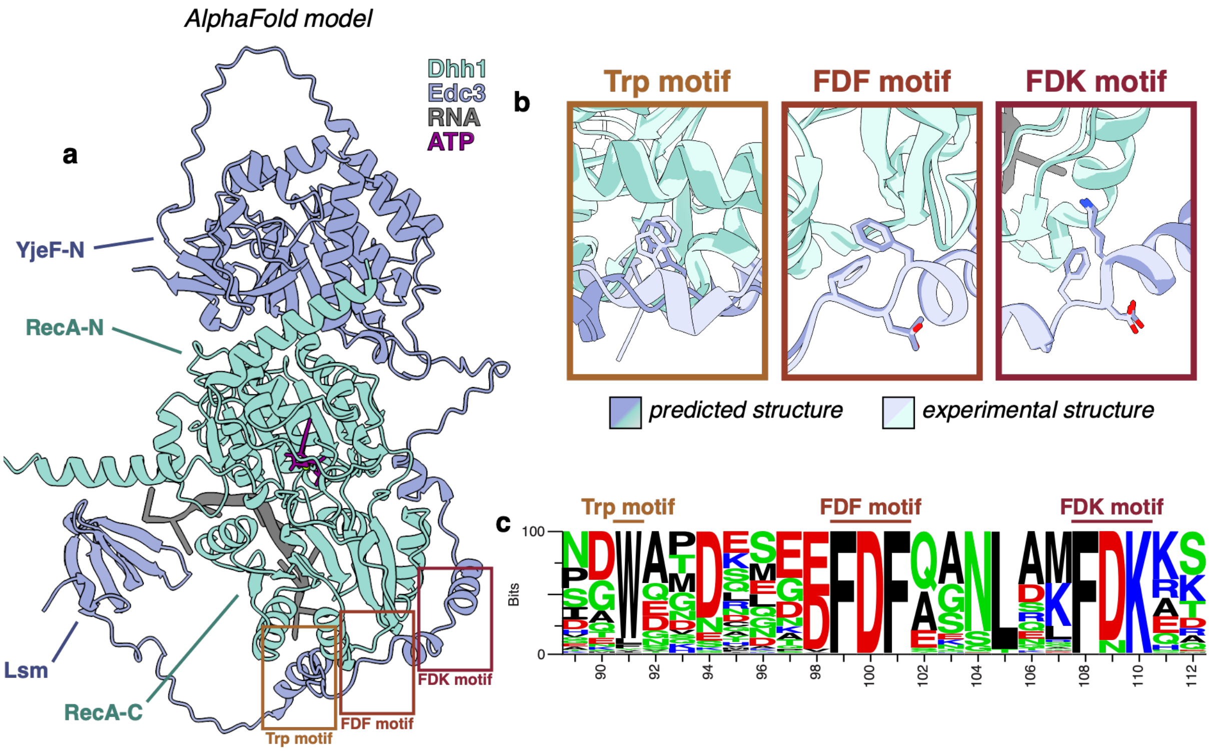


**Figure S4. Comparison of AlphaFold prediction with experimental Dhh1-Edc3 structures**
**a.** Predicted structure of Dhh1-Edc3 heterodimer, as shown in Fig. 5, with the locations of the conserved Trp, FDF, and FDK motifs indicated in boxes. **b.** Comparison of predicted interaction sites with experimental structures; for all, predicted structure is shown in dark blue and green and experimental structures are shown in light blue and green. (PDB: Trp motif: 4BRU; FDF & FDK motifs: 2WAX) **c.** Logo plot showing sequence conservation in the known Dhh1-interacting region of Edc3, with known interaction motifs marked.

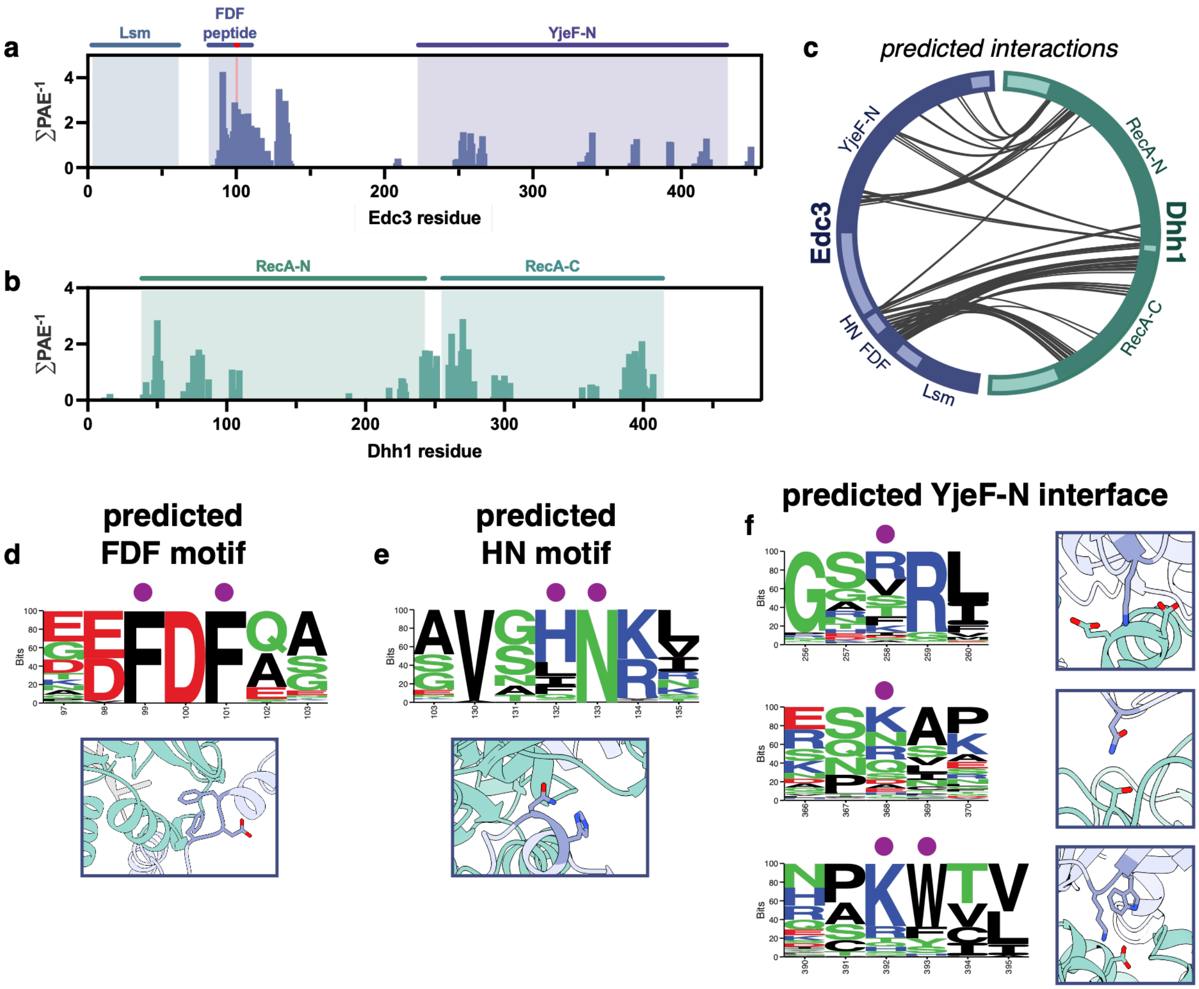


**Figure S5. Predicted Dhh1-Edc3 heterodimer includes new interaction interfaces**

**a-b.** Sites of predicted interaction between Dhh1 and Edc3 across the sequence of **(a)** Edc3 and **(b)** Dhh1, quantified as the per-residue sum of $PAE^{-1}$ for interactions under 4 Å. For each protein, domains are annotated; the FDF peptide corresponds to $Edc3^{FDF}$ construct tested experimentally (see Fig. 2e-g). **c.** Contact map of predicted Dhh1-Edc3 interactions. Lines show resides within 4 Å with PAE values < 4 Å. **d-f.** Sequence conservation and predicted interaction sites for **(d)** FDF motif, **(e)** HN motif, and **(f)** YjeF-N interface. Mutated residues are indicated with purple dots.

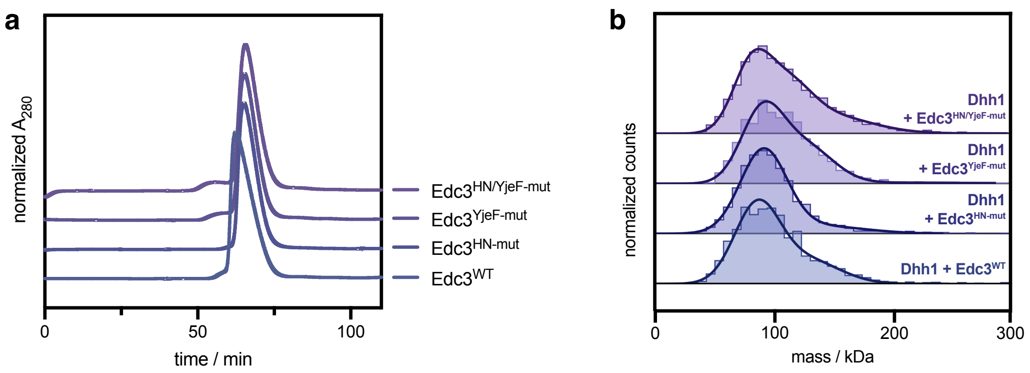


**Figure S6. Edc3 mutations do not compromise solubility or interaction with Dhh1**

**a.** Size-exclusion chromatography of $Edc3^{WT}$, $Edc3^{HN\text{-}mut}$, $Edc3^{YjeF\text{-}mut}$, and $Edc3^{HN/YjeF\text{-}mut}$. **b.** Mass photometry of binary mixtures of Dhh1 with $Edc3^{WT}$, $Edc3^{HN\text{-}mut}$, $Edc3^{YjeF\text{-}mut}$, and $Edc3^{HN/YjeF\text{-}mut}$, and $Edc3^{ADA}$. (Monomeric molecular weights: Dhh1: 55 kDa; all Edc3 variants: 49 kDa.) See **Table S9** for experimental and theoretical masses for complexes shown in panel (b).

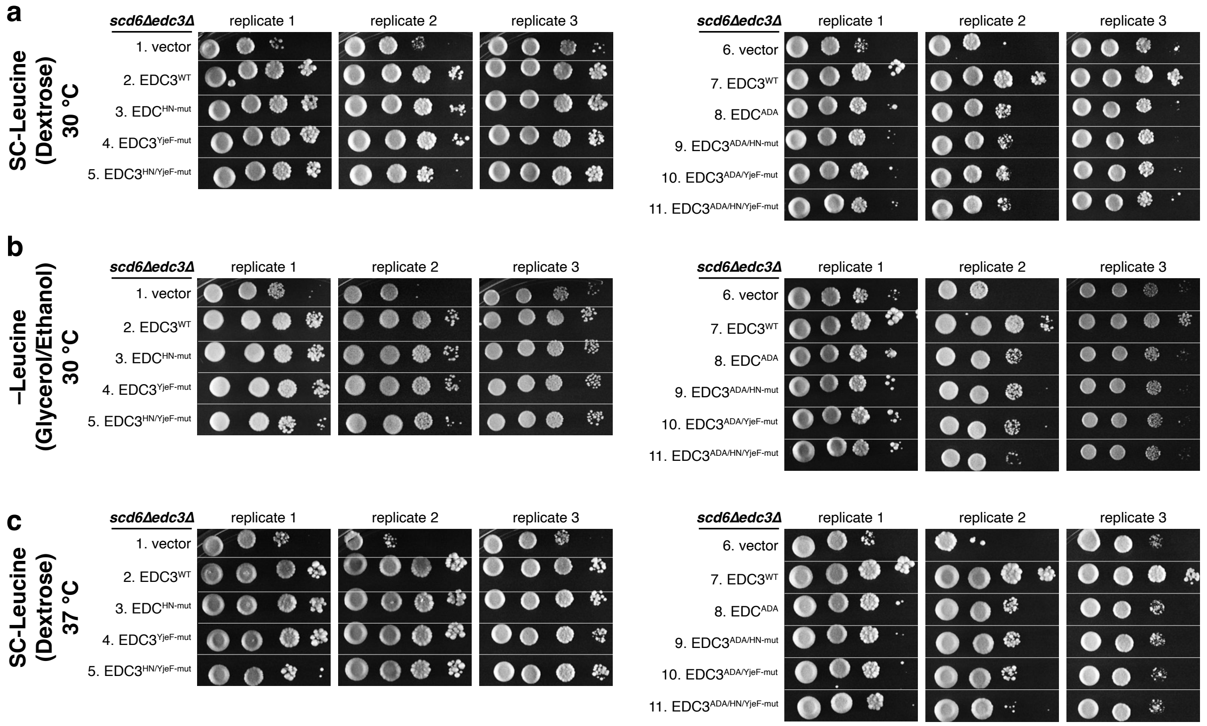


**Figure S7. Biological replicates for yeast growth from Figure 3g**

Cell growth of transformants of a *scd6ΔedcΔ S. cerevisiae* strain harboring either wild-type or mutant *EDC3* alleles, all tagged with the *myc13* epitope, on single-copy plasmids. Shown are results from three independent transformations. Cells were grown on **(a)** plates including leucine at 30 °C, **(b)** plates lacking leucine at 30 °C, and **(c)** plates including leucine at 37 °C. Spots represent 10-fold serial dilutions. Data from replicate 1 is shown in **Figure 3g**.

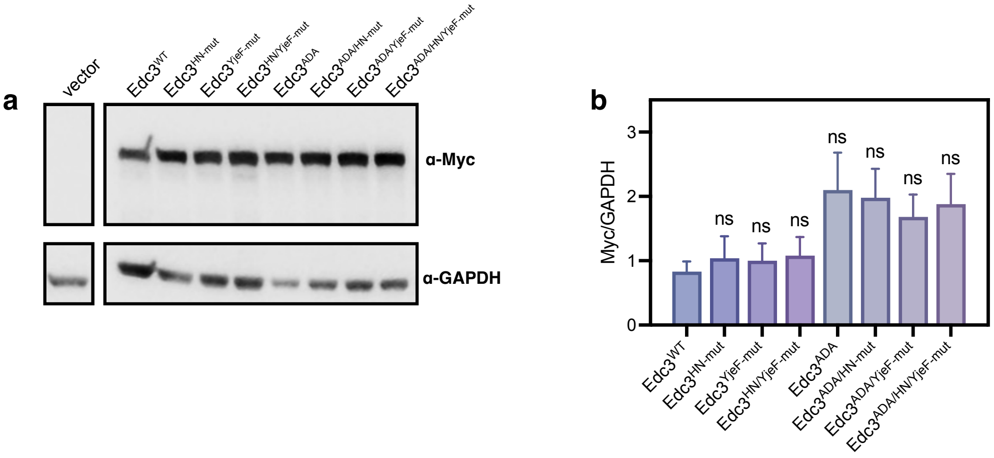


**Figure S8. Expression of wild-type and mutant *EDC3* in *S. cerevisiae***

**a.** Western blot analysis of the expression if myc13-tagged wild-type and mutant Edc3 proteins in the same transformants of the *scd6Δedc3Δ* strain analyzed for cell growth **(Fig. 3g** and **Fig. S7)**, with GAPDH used as a loading control. **b.** Quantified ratio of Myc-tagged Edc3 to GAPDH from blots of n = 3 biological replicates, with error bars showing standard deviation. (Statistical analysis: one-way ANOVA with Dunnett's multiple comparison test versus Edc3$^{WT}$, $n = 3$.) See **Table S12** for data shown in panel (b).

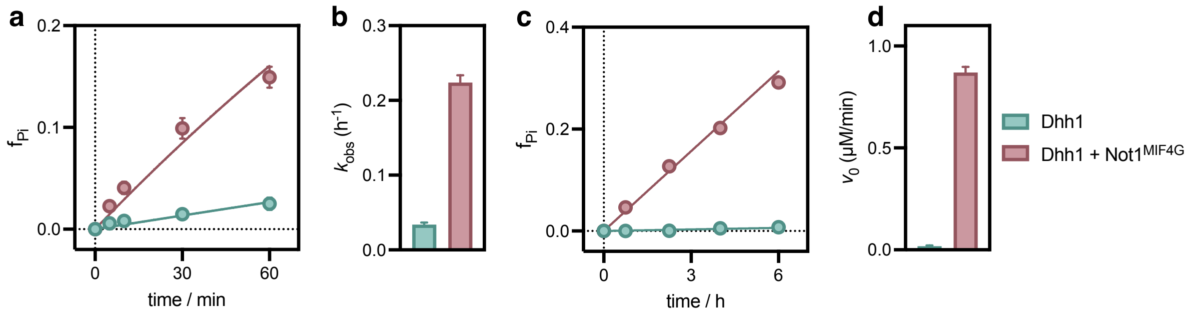


**Figure S9. Not1$^{MIF4G}$ stimulates Dhh1 ATPase activity**

ATPase activity of 5 $\mu$M Dhh1 in the absence (green) and presence (red) of 15 $\mu$M Not1$^{MIF4G}$ under **(a-b)** single-turnover and **(c-d)** multiple-turnover conditions. Total ATP concentration is 1 nM and 1 mM, respectively, and both conditions contain saturating amounts of $U_{10}$ RNA. Data points in panels (a) and (c) show mean and standard deviation of n = 3 technical replicates; error bars in panels (b) and (d) show standard deviation of fitted rates. See **Tables S3** and **S4** for single- and multiple-turnover ATPase rates shown in panels (b) and (d).

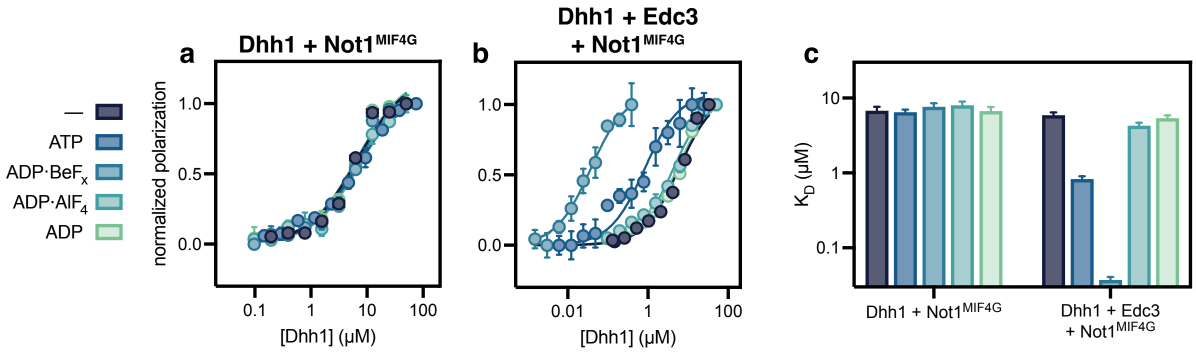


**Figure S10. Not1$^{MIF4G}$ does not affect affinity of Dhh1 for RNA in binary and ternary mixtures** Fluorescence polarization of Dhh1 binding to a fluorescein-labeled $U_{20}$ RNA in the presence of saturating amounts of **(a)** Not1$^{MIF4G}$ or **(b)** both Edc3 and Not1$^{MIF4G}$ in the absence of nucleotide or with various nucleotides and nucleotide analogs. Data points show mean and standard deviation of n = 3 technical replicates. **c.** Fitted $K_D$ values for data shown in panels (a-b), with error bars showing standard deviation. See **Table S7** for $K_D$ values shown in panel (c).

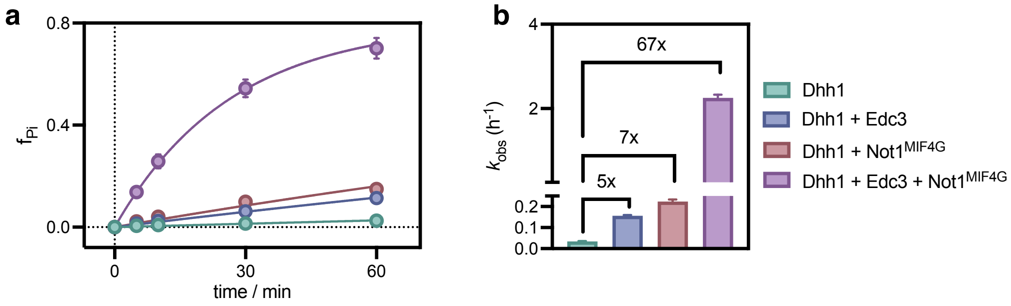


**Figure S11. Edc3 and Not1$^{MIF4G}$ cooperatively stimulate Dhh1 ATPase under single-turnover conditions**

ATPase activity of 5 $\mu$M Dhh1 alone (green) and in the presence of 15 $\mu$M Edc3 (blue), Not1$^{MIF4G}$ (red), or both Edc3 and Not1$^{MIF4G}$ (pink) under single-turnover conditions. Reactions contain saturating amounts of $U_{10}$ RNA. Data points in panel (a) show mean and standard deviation of n = 3 technical replicates; error bars in panel (b) show standard deviation of fitted rates. See **Table S3** for ATPase rates shown in panel (b).

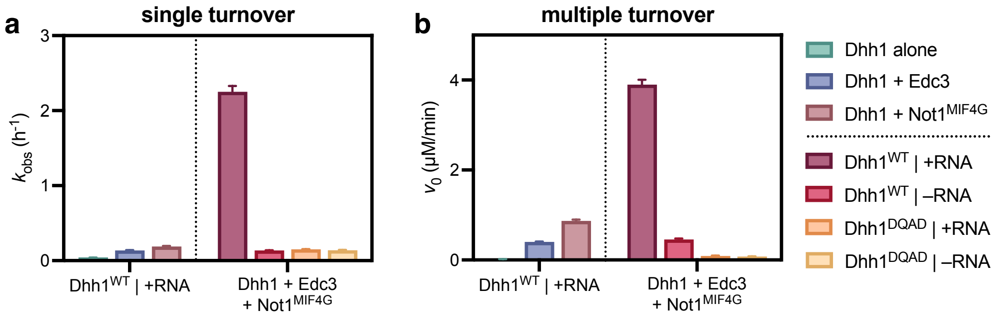


**Figure S12. Stimulation of Dhh1 by Not1 in ternary mixtures requires RNA and Walker B glutamate**

Activity of either Dhh1$^{WT}$ or Dhh1$^{DQAD}$ in the presence of both Edc3 and Not1$^{MIF4G}$ in both the presence and absence of RNA under **(a)** single-turnover and **(b)** multiple-turnover conditions. Total ATP concentration is 1 nM and 1 mM, respectively. Included for reference in each condition is the activity of Dhh1$^{WT}$ in the presence of RNA either alone (green), with Edc3 (blue), or with Not1$^{MIF4G}$ (red). Bars show fitted rate with standard deviation from n = 3 technical replicates. See **Tables S3** and **S10** for single-turnover rates shown in panel (a) and **Tables S4** and **S11** for multiple-turnover rates shown in panel (b).

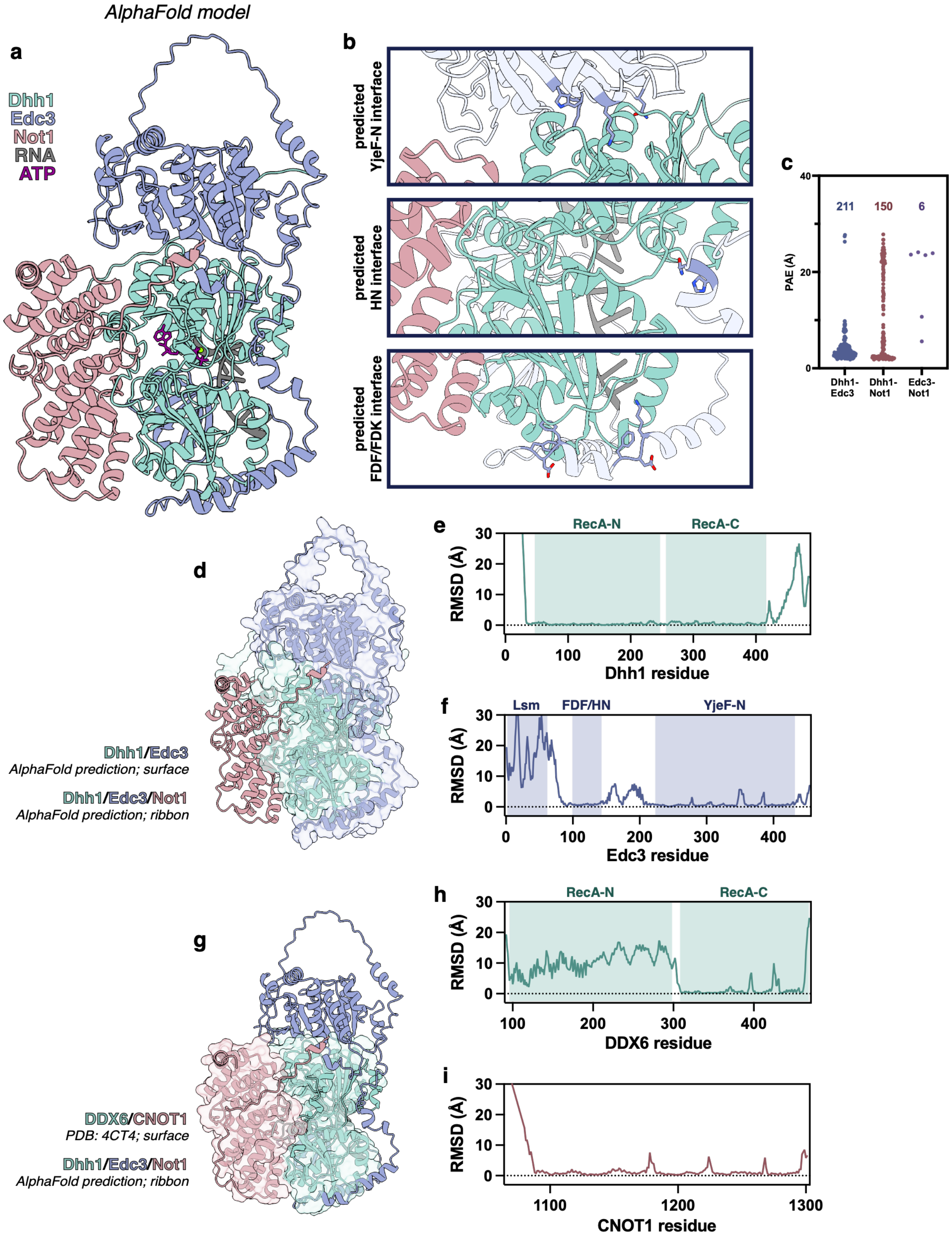


**Figure S13. Predicted structure of a heterotrimeric Dhh1-Edc3-Not1$^{MIF4G}$ complex**

**a.** Predicted structure between Dhh1 (green), Edc3 (blue), and Not1$^{MIF4G}$ (red), in the presence of $U_{10}$ RNA (grey) and ATP (purple). **b.** Close-up views of Dhh1-Edc3 interfaces in predicted

Dhh1/Edc3/Not1$^{MIF4G}$ heterotrimer shown in panel (a), demonstrating that known interfaces are unimpeded by the presence of Not1$^{MIF4G}$. Highlighted are Edc3 residues mutated at YjeF-N and HN interfaces to make Edc3$^{YjeF\text{-}mut}$ and Edc3$^{-HN\text{-}mut}$, as well as FDF and FDK motifs. **c.** Predicted aligned error (PAE) for intermolecular residue pairs within 5 Å, with the number of residue pairs shown for each pair of proteins. Low PAE indicates higher model confidence.

**d.** Comparison of predicted Dhh1/Edc3/Not1$^{MIF4G}$ heterotrimer (shown in ribbons) with predicted Dhh1/Edc3 heterodimer (shown as transparent surface). **e-f.** RMSD between structures shown in panel (d) for **(e)** Dhh1 (green) and **(f)** Edc3 (blue). **g.** Comparison of predicted Dhh1/Edc3/Not1$^{MIF4G}$ heterotrimer (shown in ribbons) with DDX6/CNOT1$^{MIF4G}$ crystal structure (PDB: 4CT4, shown as transparent surface). **h-i.** RMSD between structures shown in panel (g) for **(h)** DDX6 (green) and **(i)** CNOT1 (red). (The interdomain orientation of Dhh1 differs between the active RNA-bound predicted model and the non-RNA-bound crystal structure, leading to the uniformly high RMSD for the DDX6 RecA-N.)

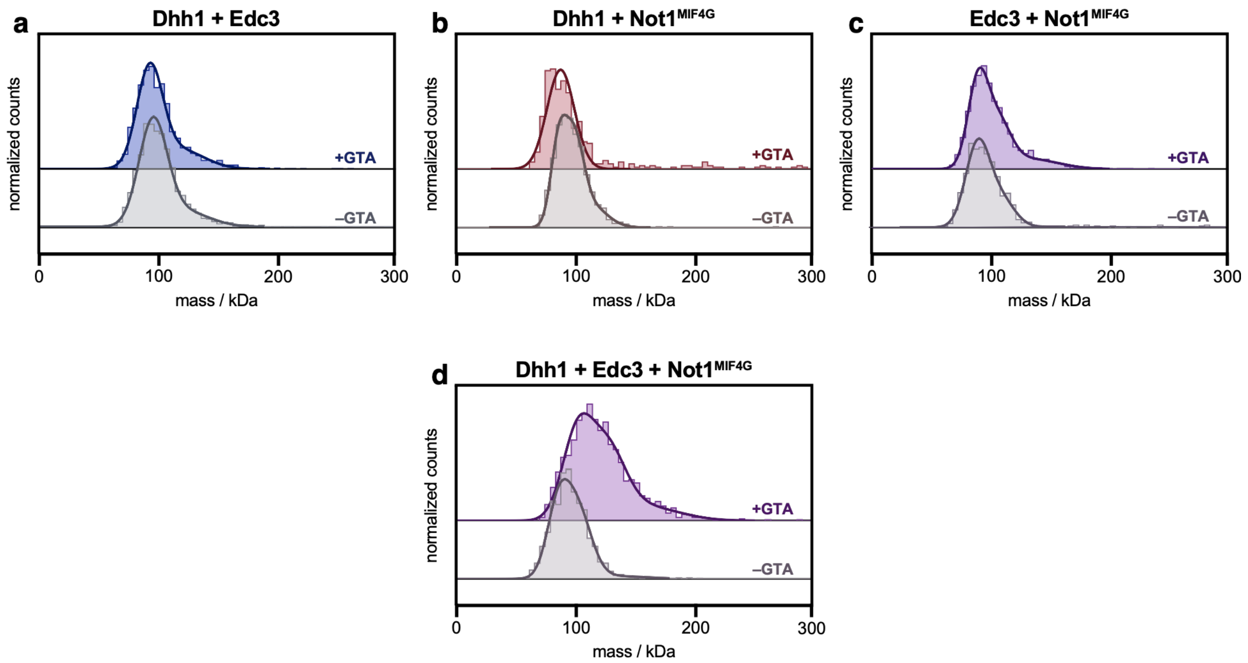


**Figure S14. Crosslinking mass photometry of binary and ternary mixtures**

Mass photometry of different combinations of Dhh1, Edc3, and $Not1^{MIF4G}$ with and without glutaraldehyde crosslinking. **a.** Binary mixture of Dhh1 and Edc3. **b.** Binary mixture of Dhh1 and $Not1^{MIF4G}$. **c.** Binary mixture of Edc3 and $Not1^{MIF4G}$. (The peak observed for this binary mixture likely corresponds to an Edc3 homodimer, with unbound $Not1^{MIF4G}$ below the detection limit of the technique.) **d.** Ternary mixture of Dhh1, Edc3, and $Not1^{MIF4G}$. See **Table S9** for experimental and theoretical masses for all complexes.

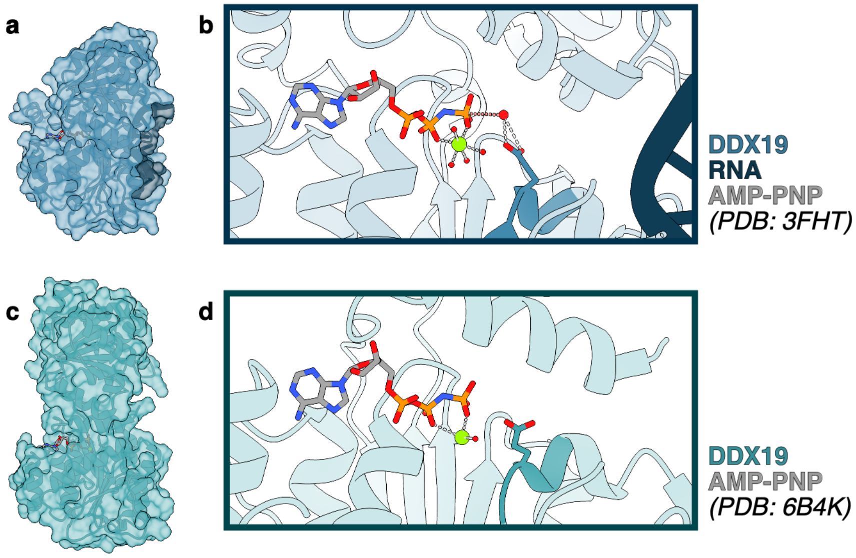


**Figure S15. Orientation of DEAD-box catalytic glutamate is sensitive to RNA**

Crystal structures of the human DBP DDX19 bound to AMP-PNP in the **(a-b)** presence and **(c-d)** absence of RNA. **a.** In the presence of RNA, DDX19 adopts a compacted tertiary structure. (PDB: 3FHT) **b.** The active site of DDX19 in the presence of RNA, with the Walker B (DEAD) motif highlighted. The glutamate of the Walker B motif forms hydrogen bonds to a water molecule positioned for hydrolysis, 3.1 Å from the AMP-PNP γ-phosphate. **c.** In the absence of RNA, DDX19 adopts an extended, open conformation. (PDB: 6B4K) **d.** The DDX19 active site in the absence of RNA. The Walker B motif is oriented away from the AMP-PNP, with no potential nucleophilic water molecules within 4 Å of the γ-phosphate.

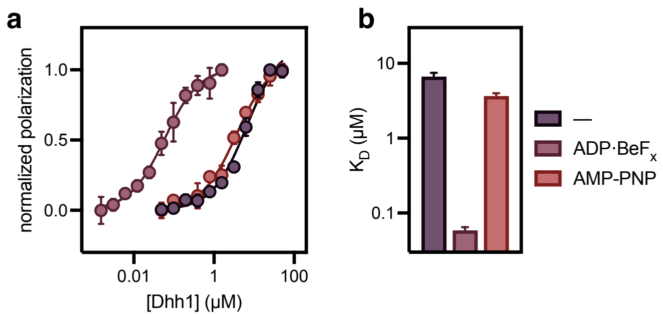


**Figure S16. Edc3 does not increase Dhh1 affinity for RNA in the presence of AMP-PNP**
**a.** Fluorescence polarization for Dhh1 binding to a fluorescein-labeled $U_{20}$ RNA in the presence of saturating amounts of Edc3 in the absence of nucleotide or with either ADP-$BeF_x$ or AMP-PNP. Data points show mean and standard deviation of n = 3 technical replicates. **b.** Fitted $K_D$ values for data shown in previous panel, with error bars showing standard deviation. Data in the absence of nucleotide and with ADP-$BeF_x$ are repeated from Figure 2 to allow for comparison. See **Table S8** for $K_D$ values shown in panel (b).

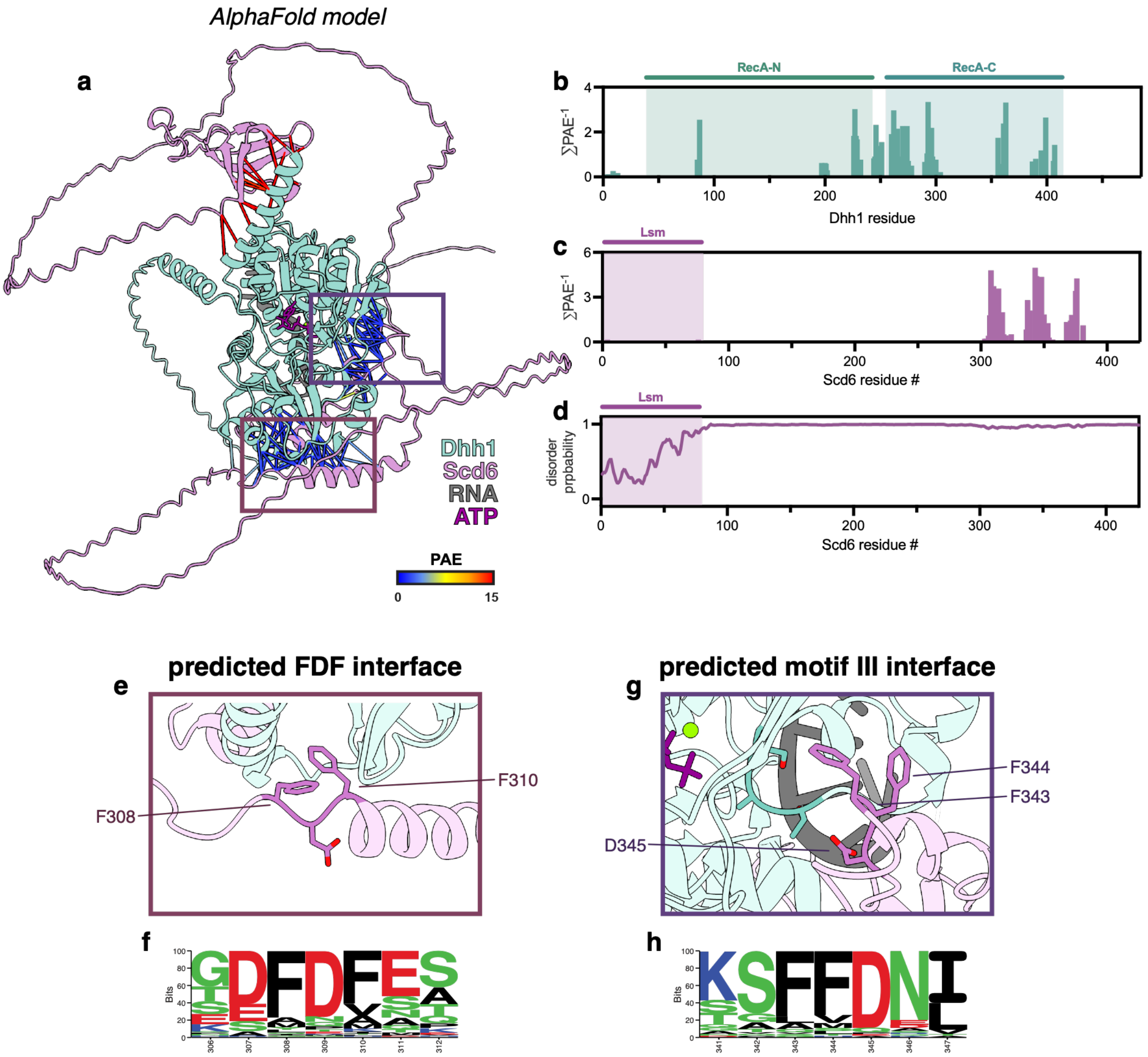


**Figure S17. Predicted Dhh1-Scd6 heterodimer includes contact analogous to HN motif**

**a.** AlphaFold 3 prediction of heterodimer between Dhh1 (green) and Scd6 (pink), in the presence of RNA (grey) and ATP (purple). Residues with predicted interprotein distances under 4 Å are shown with connecting lines, colored by Predicted Aligned Error (PAE). Boxed regions show predicted interfaces shown in greater detail in panels (e) and (g). **b-c.** Sites of predicted interaction between Dhh1 and Scd6 across the sequence of **(a)** Dhh1 and **(b)** Scd6, quantified as the per-residue sum of $PAE^{-1}$ for interactions under 4 Å. For each protein, structured domains are annotated. **d.** AIUPred disorder prediction for Scd6 (1). **e-f.** Scd6 FDF motif, showing **(e)** the predicted interaction with Dhh1 and **(f)** sequence conservation. **g-h.** Scd6 motif that interacts at similar interface to Edc3 HN motif (see Figure 3), showing **(g)** the predicted interaction with Dhh1 (Motif III of is Dhh1 colored in dark green), and **(h)** sequence conservation.

**Table S1.** Constructs used for recombinant protein expression

| protein construct | amino acid boundaries | mutations | purification tag | vector | organism |
|---|---|---|---|---|---|
| **Dhh1** | 1-485 | — | 6xHis-TEV | pETDuet | *S. pombe* |
| **Dhh1**$^{\mathbf{DQAD}}$ | 1-485 | E194Q | 6xHis-TEV | pETDuet | *S. pombe* |
| **Edc3** | 1-454 | — | 6xHis-TEV | pET-30b | *S. pombe* |
| **Edc3**$^{\mathbf{ADA}}$ | 1-454 | F99A, F101A | 6xHis-TEV | pET-30b | *S. pombe* |
| **Edc3**$^{\mathbf{\Delta Lsm}}$ | 62-454 | — | 6xHis-TEV | pET-30b | *S. pombe* |
| **Edc3**$^{\mathbf{\Delta YjeF\text{-}N}}$ | 1-192 | — | 6xHis-TEV | pET-30b | *S. pombe* |
| **Edc3**$^{\mathbf{FDF}}$ | 81-110 | — | — | pET-30b | *S. pombe* |
| **Edc3**$^{\mathbf{HN\text{-}mut}}$ | 1-454 | H132A, N133A | 6xHis-TEV | pET-30b | *S. pombe* |
| **Edc3**$^{\mathbf{YjeF\text{-}mut}}$ | 1-454 | K258E, N368A, K392E, W393A | 6xHis-TEV | pET-30b | *S. pombe* |
| **Edc3**$^{\mathbf{HN/YjeF\text{-}mut}}$ | 1-454 | H132A, N133A, K258E, N368A, K392E, W393A | 6xHis-TEV | pET-30b | *S. pombe* |
| **Not1**$^{\mathbf{MIF4G}}$ | 829-1090 | — | MBP-3C | pMAL | *S. pombe* |

**Table S2.** *EDC3* constructs used in yeast growth assays

| plasmid | mutations | epitope tag | vector | organism |
|---|---|---|---|---|
| ***EDC3***$^{WT}$ | — | 13xMyc | YCplac33 | *S. cerevisiae* |
| ***EDC3***$^{HN\text{-}mut}$ | H138A, N139A | 13xMyc | YCplac33 | *S. cerevisiae* |
| ***EDC3***$^{YjeF\text{-}mut}$ | G325Y, R331E, N336A | 13xMyc | YCplac33 | *S. cerevisiae* |
| ***EDC3***$^{HN/YjeF\text{-}mut}$ | H138A, N139A, G325Y, R331E, N336A | 13xMyc | YCplac33 | *S. cerevisiae* |
| ***EDC3***$^{ADA}$ | F105A, F107A | 13xMyc | YCplac33 | *S. cerevisiae* |
| ***EDC3***$^{ADA/HN\text{-}mut}$ | F105A, F107A, H138A, N139A | 13xMyc | YCplac33 | *S. cerevisiae* |
| ***EDC3***$^{ADA/YjeF\text{-}mut}$ | F105A, F107A, G325Y, R331E, N336A | 13xMyc | YCplac33 | *S. cerevisiae* |
| ***EDC3***$^{ADA/HN/YjeF\text{-}mut}$ | F105A, F107A, H138A, N139A, G325Y, R331E, N336A | 13xMyc | YCplac33 | *S. cerevisiae* |

**Table S3.** Single-turnover ATPase rates of Dhh1 in the presence of wild-type and mutant Edc3 and Not1$^{MIF4G}$

| | $k_{obs}$ ($h^{-1}$) | Figure(s) |
|---|---|---|
| **Dhh1** | 0.034 ± 0.003 | 2b, 2e, 3d, S1b, S9a, S11b, S12a |
| **Dhh1 + Edc3** | 0.157 ± 0.003 | S1b, S11b, S12a |
| **Dhh1 + Edc3$^{ADA}$** | 0.175 ± 0.008 | 2b |
| **Dhh1 + Edc3$^{\Delta Lsm}$** | 0.151 ± 0.007 | 2e |
| **Dhh1 + Edc3$^{\Delta YjeF\text{-}N}$** | 0.075 ± 0.008 | 2e |
| **Dhh1 + Edc3$^{FDF}$** | 0.044 ± 0.004 | 2e |
| **Dhh1 + Edc3$^{HN\text{-}mut}$** | 0.113 ± 0.006 | 3d |
| **Dhh1 + Edc3$^{YjeF\text{-}mut}$** | 0.082 ± 0.005 | 3d |
| **Dhh1 + Edc3$^{HN/YjeF\text{-}mut}$** | 0.064 ± 0.006 | 3d |
| **Dhh1 + Not1$^{MIF4G}$** | 0.224 ± 0.009 | S9b, S11b, S12a |
| **Dhh1 + Edc3 + Not1$^{MIF4G}$** | 2.25 ± 0.08 | S11b |

**Table S4.** Multiple-turnover ATPase rates of Dhh1 in the presence of wild-type and mutant Edc3 and Not1$^{MIF4G}$

| | $V_0$ (μM/min) | Figure(s) |
|---|---|---|
| **Dhh1** | 0.0244 ± 0.0015 | 2b, 2f, 3e |
| **Dhh1** | 0.017 ± 0.003 | 5b, S1d, S9d, S12b |
| **Dhh1 + Edc3** | 0.448 ± 0.012 | 2b, 2f, 3e |
| **Dhh1 + Edc3** | 0.404 ± 0.005 | 5b, S1d, S12b |
| **Dhh1 + Edc3$^{ADA}$** | 0.119 ± 0.003 | 2b |
| **Dhh1 + Edc3$^{\Delta Lsm}$** | 0.414 ± 0.009 | 2f |
| **Dhh1 + Edc3$^{\Delta YjeF\text{-}N}$** | 0.220 ± 0.014 | 2f |
| **Dhh1 + Edc3$^{FDF}$** | 0.0289 ± 0.0012 | 2f |
| **Dhh1 + Edc3$^{HN\text{-}mut}$** | 0.286 ± 0.007 | 3e |
| **Dhh1 + Edc3$^{YjeF\text{-}mut}$** | 0.167 ± 0.007 | 3e |
| **Dhh1 + Edc3$^{HN/YjeF\text{-}mut}$** | 0.144 ± 0.004 | 3e |
| **Dhh1 + Not1$^{MIF4G}$** | 0.87 ± 0.03 | 5b, S12b |
| **Dhh1 + Edc3 + Not1$^{MIF4G}$** | 3.90 ± 0.11 | 5b |

**Table S5.** Values for maximal rate ($k_{max}$) and concentration of half-maximal rate ($K_{1/2}$) from single-turnover ATPase concentration series

| | $k_{max}$ ($h^{-1}$) | $K_{1/2}$ (μM) | Figure(s) |
|---|---|---|---|
| **Dhh1** | 0.60 ± 0.05 | 90 ± 12 | 1b, 4b, 5d |
| **Dhh1 + Edc3** | 0.81 ± 0.02 | 2.0 ± 0.2 | 1b, 5d |
| **Dhh1 + Not1$^{MIF4G}$** | 4.7 ± 0.6 | 82 ± 15 | 4b, 5d |
| **Dhh1 + Edc3 + Not1$^{MIF4G}$** | 25.5 ± 1.4 | 17 ± 2 | 5d |

**Table S6.** Equilibrium dissociation constants ($K_D$) of Dhh1 for mant-ATP, in the absence and presence of Edc3

| | $K_D$ (μM) | Figure |
|---|---|---|
| **Dhh1** | 287 ± 2 | 1d |
| **Dhh1 + Edc3** | 24.7 ± 1.5 | 1d |

**Table S7.** Equilibrium dissociation constants ($K_D$) of Dhh1 for $U_{20}$ RNA in the presence of Edc3 and $Not1^{MIF4G}$ in the absence of nucleotide, or with different nucleotides and nucleotide analogs

| | **nucleotide** | **$K_D$ (μM)** | **Figure** |
|---|---|---|---|
| **Dhh1** | — | 7.0 ± 0.6 | 1e |
| | ATP | 7.5 ± 0.6 | |
| | ADP-$BeF_x$ | 7.5 ± 0.7 | |
| | ADP-$AlF_4$ | 7.0 ± 0.6 | |
| | ADP | 9.1 ± 0.8 | |
| **Dhh1 + Edc3** | — | 6.6 ± 0.8 | 1e |
| | ATP | 0.44 ± 0.07 | |
| | ADP-$BeF_x$ | 0.059 ± 0.006 | |
| | ADP-$AlF_4$ | 7.6 ± 1.4 | |
| | ADP | 7.0 ± 0.9 | |
| **Dhh1 + $Not1^{MIF4G}$** | — | 6.8 ± 0.9 | S10c |
| | ATP | 6.4 ± 0.5 | |
| | ADP-$BeF_x$ | 7.7 ± 0.8 | |
| | ADP-$AlF_4$ | 8.0 ± 1.0 | |
| | ADP | 6.7 ± 0.9 | |
| **Dhh1 + Edc3 + $Not1^{MIF4G}$** | — | 5.9 ± 0.5 | S10c |
| | ATP | 0.83 ± 0.08 | |
| | ADP-$BeF_x$ | 0.037 ± 0.003 | |
| | ADP-$AlF_4$ | 4.3 ± 0.4 | |
| | ADP | 5.3 ± 0.5 | |

**Table S8.** Equilibrium dissociation constants ($K_D$) of Dhh1 for $U_{20}$ RNA in the presence of wild-type and mutant Edc3 (unless otherwise noted, all data are in the presence of ADP-BeF$_x$)

| | $K_D$ ($\mu$M) | Figure(s) |
|---|---|---|
| **Dhh1** | 6.3 ± 0.4 | 2c, 2g, 3f, S15b |
| **Dhh1 + Edc3** | 0.059 ± 0.006 | 2c, 2g, 3f, S15b |
| **Dhh1 + Edc3$^{ADA}$** | 6.9 ± 0.5 | 2c |
| **Dhh1 + Edc3$^{\Delta Lsm}$** | 0.061 ± 0.004 | 2g |
| **Dhh1 + Edc3$^{\Delta YjeF\text{-}N}$** | 4.5 ± 0.3 | 2g, 3f |
| **Dhh1 + Edc3$^{FDF}$** | 4.7 ± 0.3 | 2g |
| **Dhh1 + Edc3$^{HN\text{-}mut}$** | 0.130 ± 0.009 | 3f |
| **Dhh1 + Edc3$^{YjeF\text{-}mut}$** | 3.6 ± 0.2 | 3f |
| **Dhh1 + Edc3$^{HN/YjeF\text{-}mut}$** | 4.7 ± 0.3 | 3f |
| **Dhh1 + Edc3** *(AMP-PNP)* | 3.6 ± 0.3 | S16b |

**Table S9.** Experimental masses for individual proteins and protein complexes as determined by mass photometry

| | experimental mass (kDa) | theoretical mass (kDa) | Figure(s) |
|---|---|---|---|
| **Dhh1** | 49 ± 13 | Dhh1: 54.9 | 2h, 5e |
| **Dhh1** | 66 ±18 | Dhh1: 54.9 | S2b |
| **Edc3** | 95 ± 13 | Edc3: 49.4<br>Edc3 dimer: 98.7 | 2h |
| **Dhh1 + Edc3** | 98 ± 18 | Dhh1: 54.9<br>Edc3: 49.4<br>Dhh1-Edc3 heterodimer: 104.2 | 2h, 5e, S2a, S6b |
| **Dhh1 + Edc3** | 99 ± 25 | Dhh1: 54.9<br>Edc3: 49.4<br>Dhh1-Edc3 heterodimer: 104.2 | S2b |
| **Dhh1 + Edc3** | 99 ± 18 | Dhh1: 54.9<br>Edc3: 49.4<br>Dhh1-Edc3 heterodimer: 104.2 | S14a |
| **Dhh1 + Edc3** (+ glutaraldehyde) | 97 ± 18 | Dhh1: 54.9<br>Edc3: 49.4<br>Dhh1-Edc3 heterodimer: 104.2 | S14a |
| **Dhh1 + Edc3$^{ADA}$** (150 mM NaCl) | 97 ± 20 | Dhh1: 54.9<br>Edc3$^{ADA}$: 49.2<br>Dhh1-Edc3$^{ADA}$ heterodimer: 104.0 | S2a |
| **Dhh1 + Edc3$^{WT}$** (500 mM NaCl) | 101 ± 30 | Dhh1: 54.9<br>Edc3: 49.4<br>Dhh1-Edc3 heterodimer: 104.2 | S2a |
| **Dhh1 + Edc3$^{ADA}$** (500 mM NaCl) | 67 ± 17<br>86 ± 24 | Dhh1: 54.9<br>Edc3$^{ADA}$: 49.2<br>Edc3$^{ADA}$ homodimer: 98.3<br>Dhh1-Edc3$^{ADA}$ heterodimer: 104.0 | S2a |
| **Dhh1 + Edc3$^{\Delta YjeF\text{-}N}$** | 73 ± 17 | Dhh1: 54.9<br>Edc3$^{\Delta YjeF\text{-}N}$: 21.3<br>Dhh1-Edc3$^{\Delta YjeF\text{-}N}$ heterodimer: 76.2 | S2b |
| **Dhh1 + Edc3$^{HN\text{-}mut}$** | 98 ± 18 | Dhh1: 54.9<br>Edc3$^{HN\text{-}mut}$: 49.3<br>Dhh1-Edc3$^{HN\text{-}mut}$ heterodimer: 104.1 | S6b |
| **Dhh1 + Edc3$^{YjeF\text{-}mut}$** | 103 ± 2 | Dhh1: 54.9<br>Edc3$^{YjeF\text{-}mut}$: 49.2<br>Dhh1-Edc3$^{YjeF\text{-}mut}$ heterodimer: 104.1 | S6b |
| **Dhh1 + Edc3$^{HN/YjeF\text{-}mut}$** | 103 ± 24 | Dhh1: 54.9<br>Edc3$^{YjeF\text{-}mut}$: 49.1<br>Dhh1-Edc3$^{YjeF\text{-}mut}$ heterodimer: 104.0 | S6b |
| **Dhh1 + Not1$^{MIF4G}$** | 89 ± 14 | Dhh1: 54.9<br>Not1$^{MIF4G}$: 30.3<br>Dhh1-Not1$^{MIF4G}$ heterodimer: 85.2 | 5e |

| | | | |
|---|---|---|---|
| **Dhh1 + Not1$^{MIF4G}$** | 90 ± 13 | Dhh1: 54.9<br>Not1$^{MIF4G}$: 30.3<br>Dhh1-Not1$^{MIF4G}$ heterodimer: 85.2 | S14b |
| **Dhh1 + Not1$^{MIF4G}$**<br>(+ glutaraldehyde) | 88 ± 12 | Dhh1: 54.9<br>Not1$^{MIF4G}$: 30.3<br>Dhh1-Not1$^{MIF4G}$ heterodimer: 85.2 | S14b |
| **Edc3 + Not1$^{MIF4G}$** | 96 ± 14 | Edc3: 49.4<br>Edc3 homodimer: 98.7<br>Not1$^{MIF4G}$: 30.3 | S14c |
| **Edc3 + Not1$^{MIF4G}$**<br>(+ glutaraldehyde) | 98 ± 20 | Edc3: 49.4<br>Edc3 homodimer: 98.7<br>Not1$^{MIF4G}$: 30.3 | S14c |
| **Dhh1 + Edc3 + Not1$^{MIF4G}$** | 95 ± 22 | Dhh1: 54.9<br>Edc3: 49.4<br>Not1$^{MIF4G}$: 30.3<br>Dhh1-Edc3 heterodimer: 104.2<br>Dhh1-Edc3- Not1$^{MIF4G}$: heterotrimer: 134.5 | 5e |
| **Dhh1 + Edc3 + Not1$^{MIF4G}$** | 95 ± 17 | Dhh1: 54.9<br>Edc3: 49.4<br>Not1$^{MIF4G}$: 30.3<br>Dhh1-Edc3 heterodimer: 104.2<br>Dhh1-Edc3- Not1$^{MIF4G}$: heterotrimer: 134.5 | S14d |
| **Dhh1 + Edc3 + Not1$^{MIF4G}$**<br>(+ glutaraldehyde) | 127 ± 26 | Dhh1: 54.9<br>Edc3: 49.4<br>Not1$^{MIF4G}$: 30.3<br>Dhh1-Edc3 heterodimer: 104.2<br>Dhh1-Edc3- Not1$^{MIF4G}$: heterotrimer: 134.5 | 5e |
| **Dhh1 + Edc3 + Not1$^{MIF4G}$**<br>(+ glutaraldehyde) | 118 ± 27 | Dhh1: 54.9<br>Edc3: 49.4<br>Not1$^{MIF4G}$: 30.3<br>Dhh1-Edc3 heterodimer: 104.2<br>Dhh1-Edc3- Not1$^{MIF4G}$: heterotrimer: 134.5 | S14d |

**Table S10.** Single-turnover ATPase rates of $Dhh1^{WT}$ and $Dhh1^{DQAD}$ in the presence of Edc3 and $Not1^{MIF4G}$ in either the presence or absence of RNA

| | | | $k_{obs}$ ($h^{-1}$) | Figure |
|---|---|---|---|---|
| **Dhh1** | $Dhh1^{WT}$ | +RNA | 0.036 ± 0.003 | 4c |
| | | –RNA | 0.033 ± 0.003 | 4c |
| | $Dhh1^{DQAD}$ | +RNA | 0.035 ± 0.003 | 4c |
| | | –RNA | 0.033 ± 0.002 | 4c |
| **Dhh1 + $Not1^{MIF4G}$** | $Dhh1^{WT}$ | +RNA | 0.188 ± 0.009 | 4c |
| | | –RNA | 0.0339 ± 0.0016 | 4c |
| | $Dhh1^{DQAD}$ | +RNA | 0.039 ± 0.003 | 4c |
| | | –RNA | 0.0344 ± 0.0019 | 4c |
| **Dhh1 + Edc3** | $Dhh1^{WT}$ | +RNA | 0.136 ± 0.004 | 4c |
| | | –RNA | 0.128 ± 0.005 | 4c |
| | $Dhh1^{DQAD}$ | +RNA | 0.143 ± 0.004 | 4c |
| | | –RNA | 0.132 ± 0.004 | 4c |
| **Dhh1 + Edc3 + $Not1^{MIF4G}$** | $Dhh1^{WT}$ | +RNA | 2.25 ± 0.08 | S12a |
| | | –RNA | 0.136 ± 0.003 | S12a |
| | $Dhh1^{DQAD}$ | +RNA | 0.148 ± 0.004 | S12a |
| | | –RNA | 0.139 ± 0.005 | S12a |

**Table S11.** Multiple-turnover ATPase rates of Dhh1$^{WT}$ and Dhh1$^{DQAD}$ in the presence of Edc3 and Not1$^{MIF4G}$ in either the presence or absence of RNA

| | | | $V_0$ (μM/min) | Figure(s) |
|---|---|---|---|---|
| **Dhh1** | Dhh1$^{WT}$ | +RNA | 0.017 ± 0.003 | 4d |
| | | –RNA | 0.0015 ± 0.0003 | 4d |
| | Dhh1$^{DQAD}$ | +RNA | 0.0042 ± 0.0011 | 4d |
| | | –RNA | 0.0017 ± 0.0003 | 4d |
| **Dhh1 + Not1$^{MIF4G}$** | Dhh1$^{WT}$ | +RNA | 0.87 ± 0.03 | 4d |
| | | –RNA | 0.044 ± 0.002 | 4d |
| | Dhh1$^{DQAD}$ | +RNA | 0.043 ± 0.003 | 4d |
| | | –RNA | 0.032 ± 0.002 | 4d |
| **Dhh1 + Edc3** | Dhh1$^{WT}$ | +RNA | 0.404 ± 0.005 | 4d |
| | | –RNA | 0.248 ± 0.005 | 4d |
| | Dhh1$^{DQAD}$ | +RNA | 0.051 ± 0.003 | 4d |
| | | –RNA | 0.040 ± 0.002 | 4d |
| **Dhh1 + Edc3 + Not1$^{MIF4G}$** | Dhh1$^{WT}$ | +RNA | 3.89 ± 0.11 | S12b |
| | | –RNA | 0.458 ± 0.017 | S12b |
| | Dhh1$^{DQAD}$ | +RNA | 0.091 ± 0.006 | S12b |
| | | –RNA | 0.079 ± 0.004 | S12b |

**Table S12.** Western blot quantification of Myc-tagged Edc3, normalized to GAPDH loading control (related to Fig. S8)

| | Myc/GAPDH |
|---|---|
| **Edc3$^{WT}$** | 0.83 ± 0.16 |
| **Edc3$^{HN\text{-}mut}$** | 1.0 ± 0.3 |
| **Edc3$^{YjeF\text{-}mut}$** | 1.0 ± 0.3 |
| **Edc3$^{HN/YjeF\text{-}mut}$** | 1.1 ± 0.3 |
| **Edc3$^{ADA}$** | 2.1 ± 0.6 |
| **Edc3$^{ADA/HN\text{-}mut}$** | 2.0 ± 0.5 |
| **Edc3$^{ADA/YjeF\text{-}mut}$** | 1.7 ± 0.4 |
| **Edc3$^{ADA/HN/YjeF\text{-}mut}$** | 1.9 ± 0.5 |

### SUPPLEMENTAL METHODS

#### Sequence Conservation Analysis

Sequence conservation of Edc3 and Scd6 was generated using ConSurf (2); per-residue conservation was visualized using Seq2Logo (3).

#### Western Blotting

Samples from *scd6Δedc3Δ* transformants used in cell growth spotting assays were prepared by trichloroacetic acid extraction as previously described (4), and immunoblot analysis was performed as previously described (5). Steady-state Edc3 expression was probed using anti-Myc antibody with GAPDH used as a loading control. Densitometric analysis of western blot bands was performed using ImageJ. The signal intensity of each Myc-tagged protein was formalized to its respective GAPDH loading control.
